# An Engineered 3D Co-culture Model of Primary Macrophages and Patient-Derived Tumour Cells to Explore Cellular Responses in the Graded Hypoxic Microenvironment of Pancreatic Cancer

**DOI:** 10.1101/2023.07.13.548899

**Authors:** Ileana L. Co, Chengxin Yu, Sara Lamorte, M. Teresa Ciudad, Natalie Landon-Brace, Jose L. Cadavid, Ziting Xia, Aleksandra Fomina, Michelle Nurse, Tracy L. McGaha, Kieran R. Campbell, Alison P. McGuigan

## Abstract

In pancreatic ductal adenocarcinoma (PDAC), tumour associated macrophages (TAMs) are a heterogeneous immune cell population that interact with cancer cells to promote malignancy, chemo-resistance, and immunosuppression. Aside from TAMs, hypoxia is a prominent feature of PDAC that can rewire cells to survive and enhance malignancy in the tumour microenvironment (TME). Deciphering the interactions between macrophages, cancer cells and hypoxia could lead to the development of effective immune-targeted therapies for PDAC. However, there are only a few models that physiologically recapitulate the PDAC TME and allow for meaningful interrogation of cancer-immune cell interactions in hypoxia. Here, we develop a model of primary macrophages and PDAC patient organoid-derived cells by adapting TRACER, a paper-based, engineered 3D model that allows snapshot analysis of cellular response in hypoxia. In this study, we establish a direct co-culture method of primary macrophages and PDAC organoid cells in TRACER and demonstrate that TRACER co-cultures generate hypoxic gradients and show expected phenotypic responses to this hypoxic gradient. Moreover, we report for the first time in a human *in vitro* model that hypoxic macrophages exert a graded chemoprotective effect on gemcitabine-treated PDAC organoid cells, and that interactions between cancer cells and macrophages from the inner layers of TRACER indirectly attenuate the inflammatory response of donor-derived T-cells. Overall, the TRACER co-culture system is a novel, fully human 3D *in vitro* cancer-immune model for evaluating the response of macrophages and cancer cells in a hypoxic gradient.

## 1. Introduction

Immune-targeted therapies are promising treatments that aim to modulate the immune system to combat cancer. In the clinic, they have shown success in previously hard-to-treat cancers like melanoma and non-small cell lung carcinomas^1^. However, for many solid cancers like pancreatic ductal adenocarcinoma (PDAC), immunotherapies such as checkpoint blockade therapy, still fail. One key reason why immunotherapies might be ineffective in PDAC is due to their highly heterogeneous and dysregulated tumour microenvironment (TME). Specifically, the PDAC TME is characterized by an extremely fibrotic stroma, an immunosuppressive immune infiltrate, a highly hypoxic microenvironment, and is comprised of various cellular and acellular components that dynamically interact with each other to orchestrate a pro-tumorigenic environment^2–4^. Of the interactions in the TME, cancer-immune interactions are significant due to their role in modulating the TME to promote malignancy and their potential to be targeted as immunotherapies^2^. Specifically, tumour associated macrophages (TAMs) are a class of highly plastic and heterogeneous macrophages that can be resident in the tumour tissue or recruited to the TME by cancer cells and can display a mixture of pro and anti-tumorigenic behaviours depending on their microenvironment. In PDAC, TAMs are one of the most abundant immune cells, comprising around 10-20% of the total tumour mass, and have been correlated with both poorer overall survival and the more aggressive basal PDAC subtype^5–7^. Notably, the highly hypoxic microenvironment of PDAC has been shown to reprogram TAMs to impair cytotoxic T-cell effector function, remodel the ECM, and promote metastasis and angiogenesis, suggesting that hypoxia is an important driver of pro-tumorigenic TAM phenotypes^8–11^. Overall, TAMs and their interactions with microenvironmental factors in the TME, such as hypoxia, play influential roles in orchestrating disease progression. Investigating these interactions more deeply could help decipher why immune-targeted therapies are not successful and could lead to the development of more effective therapeutic strategies for PDAC.

Despite the importance of understanding TAM-tumour interactions in hypoxia, modeling these interactions are typically not prioritized; often, studies are focused solely on the tumour cells and not on microenvironmental interactions. This is likely because exploring these intersecting parameters is challenging to do using current cancer models. Some of these models include oversimplified cell line-based 2D models which do not model the complex interactions in the TME^12^, *in vivo* mouse models which capture the complexities of the TME but suffer from species-mismatch from humans^13,14^, and patient tissue which are physiologically relevant but are challenging to obtain and use for interrogating transient features like hypoxia. Recently, 3D models such as patient-derived organoids (PDOs) have emerged as systems that better recapitulate patient tissue by growing cells in an ECM. PDOs have successfully been created for various cancers like PDAC, and have importantly demonstrated morphological, genetic, and phenotypic similarities to their tumour of origin^15–17^. While several studies have improved the complexity of PDOs by integrating important cellular components such as cancer associated fibroblasts (CAFs)^18–21^, very few attempts have been reported to integrate primary macrophages into organoid cultures to explore cancer-immune cell interactions^22–26;^ similarly, only a handful of models attempt to explore intersecting parameters between hypoxia, cancer cells, and immune cells^27–31^.

To address this gap, we set out to develop an engineered 3D cancer-immune model of PDAC by adapting a platform developed in our group called the Tissue Roll for Analysis of Cellular and Environment Response (TRACER). TRACER is an engineered, paper-supported 3D model wherein cancer cells embedded in a matrix are infiltrated into a cellulose scaffold. This scaffold is rolled around an oxygen impermeable mandrel to create a layered-stack of cells that can be rapidly unrolled to allow for layer-specific analysis. In layers closer to the centre of the roll and progressively further from the media, cells have less access to nutrients and oxygen due to cellular consumption by the outer layers, forming a cell-generated gradient of hypoxia and metabolites^32–34^. Notably, our group has adapted the original TRACER model to be compatible for culturing primary cells like PDAC PDOs and demonstrated its use in setting-up co-cultures with pancreatic stellate cells to investigate phenotypic changes due to hypoxic gradients^21,35^. In this study, we co-cultured peripheral blood mononuclear cell (PBMC)-derived macrophages directly with organoid-derived cells (ODCs) from PDAC PDOs using the previously developed TRACER system. We first investigated the feasibility of culturing primary macrophages in TRACER and optimized co-culture methods and media conditions. Then, we show that ODCs and macrophages experienced hypoxic gradients in TRACER and display expected phenotypic responses to these gradients. To demonstrate the value of our model in investigating functional and phenotypic cellular changes, we developed assays to evaluate the response of the TRACER co-culture to the standard-of-care chemotherapy gemcitabine, as well as the response of T-cells to macrophage-tumour interactions in hypoxia. Overall, in this study, we describe a novel, engineered, fully human primary 3D model of pancreatic cancer and macrophages and investigate the rewiring of macrophage-cancer cell response in a hypoxic gradient. Having demonstrated the use and functionalities of this unique cancer-immune model, we envision that our model will enable the interrogation of the intricate interactions between macrophages and cancer cells and aid in developing more effective therapeutics for solid cancers like PDAC.

## 2. Materials and Methods

### Isolation and freezing of PBMCs from donor-derived buffy coats

As previously described^36^, PBMCs were extracted from buffy coats obtained from healthy human donors supplied by the Canadian Blood Services Blood4Research Program (CBSREB# 2018.020) using the Ficoll-paque ® density gradient centrifugation method. Buffy coats were diluted with phosphate buffered saline (PBS) without Calcium and Magnesium (Wisent Bio Products) in a 1:3 ratio, then carefully distributed into tubes containing a layer of Ficoll-paque ™ (Cytiva). The tubes were then centrifuged at 500 G at room temperature for 30 minutes. After centrifugation, the buffy coat components were separated by density into a new tube. Then, the red blood cells were lysed using an Ammonium Chloride based buffer^37^, and the PBMC-layer was extracted. Typically, freshly isolated PBMCs were resuspended in a 1:1 ratio of 5% Fetal Bovine Serum (FBS) in PBS and 20% dimethyl sulfoxide (DMSO) in FBS for cryopreservation in liquid nitrogen. All buffy coats used in this study were approved for use by the University of Toronto Research Ethics Board (Protocol no. 25841).

### CD14-monocyte positive selection and macrophage differentiation

To obtain CD14^+^ monocytes, PBMCs were quickly thawed and resuspended in ice-cold RPMI-1640 media (Gibco) supplemented with 10% FBS and 1% Penicillin-Streptomycin (RPMI-1640 (complete)) to help maintain cell viability. PBMCs were then labeled with CD14 antibodies conjugated to magnetic beads (Miltenyi Biotec). Magnetically labeled PBMCs were then passed through an LS column (Miltenyi Biotec) to allow for positive selection of CD14^+^ monocytes, as per the manufacturer’s protocol. Freshly selected CD14^+^ monocytes were counted using a haemocytometer and plated in tissue-culture treated dishes with RPMI-1640 (complete) for differentiation into macrophages. To differentiate freshly isolated CD14^+^ monocytes to M0 macrophages, 50 ng/mL M-CSF (Peprotech) was added to the media and cells were left in the incubator at 37°C and 5% CO2 for 5-7 days. All cells tested negative for mycoplasma during monthly testing.

### Macrophage seeding and activation in 2D, gel, and single-layer TRACER scaffolds

To determine the feasibility of culturing macrophages in TRACER scaffolds, M0 (unstimulated) macrophages were seeded in either 2D tissue culture plates, embedded in a gel blend, or seeded onto TRACER scaffolds. For the 2D configuration, macrophages were seeded at a density of 800,000 cells/mL in a 6-well plate; For the gel configuration, 200,000 macrophages were seeded as 15 µL domes in a blend of 75% neutralized Type I Bovine Collagen (Pur Col 3mg/mL) and 25% Matrigel®; For TRACER scaffolds, 200,000 macrophages were resuspended in 15 µL of the gel blend. Subsequently, 5 µL of the cell/gel suspension was spotted onto a polydimethylsiloxane (PDMS) (Dow Chemicals) seeding device (as manufactured previously^32–34)^ and immediately overlaid with a single-layer TRACER scaffold. The seeding device was then placed in a humidification chamber at 37°C for 45 minutes to allow for gelation then transferred into 48-well plates with RPMI-1640 (complete) media. For all three configurations, M0 macrophages were cultured for 24 hours in 10 ng/mL M-CSF in an incubator to maintain cell viability. To evaluate the plasticity of macrophages, M0 macrophages in 2D and single-layer TRACER scaffolds were additionally stimulated for another 24 hours with 50 ng/mL IFNγ (Peprotech) and 10 ng/mL lipopolysaccharide (LPS) (Gibco) for activation to pro-inflammatory phenotypes, or 20 ng/mL IL-4 (Peprotech) and 20 ng/mL IL-13 (Peprotech) for anti-inflammatory phenotypes, as performed previously^36,38–40^.

### Gene expression analysis of macrophage monocultures via qPCR

To analyze gene expression of macrophages in 2D, gel and TRACER scaffolds, the RNeasy mini kit (Qiagen) was used to extract the RNA from the cells. For each configuration, cells were washed once with PBS to remove residual media and then lysed with 500 µL RLT buffer (Qiagen) with 5 µL of β-mercaptoethanol (Sigma). Specifically for the gel configuration, gel domes were scraped off the plate with a micropipette tip and transferred to 1.5mL tubes for lysis; for the single-layer TRACER configuration, scaffolds were transferred directly to 1.5mL tubes for lysis. For both gel and single-layer TRACERs, the tubes were vortexed for 5-10 mins to facilitate cell lysis. The remainder of the RNA extraction steps were then completed following the manufacturer’s instructions. The extracted RNA was quantified using a NanoDrop Spectrophotometer (ThermoFisher Scientific) and immediately reverse transcribed to cDNA using the qScript cDNA SuperMix (QuantaBio). 8 ng of cDNA was used for qPCR, where gene-specific primers described in **Table S2** were amplified and detected using the PerfeCTa SYBR® Green FastMix (QuantaBio). The qPCR was performed using a CFX-384 thermocycler (BioRad), while data were acquired using Bio-Rad CFX Manager 3.1. Gene expression was normalized to the housekeeping genes GAPDH and 18S and the 2^−ΔΔCt^ method was used to determine relative gene expression.

### Cell culture of PDAC PDOs

The PPTO.46 PDAC organoid model was obtained from the Princess Margaret Living Biobank, Toronto, Ontario, Canada. In this study, PDOs were dissociated into single cells termed organoid-derived cells (ODCs) for culture in TRACER, as previously described^35^. PPTO.46 cells were maintained in organoid complete growth media (CGM) which was composed of Advanced DMEM/F-12 (Gibco) supplemented with 2 mM GlutaMAX, 10 mM HEPES (Gibco), 1% penicillin/streptomycin, 1X W21 supplement (Wisent), 1.25 mM N-acetyl-l-cysteine, 10 nM Gastrin I (1-14) (Sigma-Aldrich), 50 ng/mL recombinant human EGF, 100 ng/mL recombinant human noggin, 100 ng/mL recombinant human FGF-10 (Peprotech), 0.5 μM A83-01 (Tocris Biosciences), 10 μM Y-27632 (Selleck Chemicals), 10 mM nicotinamide (Sigma-Aldrich), 20% v/v Wnt-3a conditioned media, and 30% v/v human R-spondin1 conditioned media (Princess Margaret Living Biobank, Toronto, Canada). PPTO.46 cells were cultured in 48-well tissue-culture treated plates in 50 μL domes of Growth Factor Reduced Phenol Red-Free Matrigel® Matrix (Corning Life Sciences) with 500 μL of organoid CGM. PPTO.46 cells were passaged once per week (1:8 split ratio) and typically used for seeding after 6-8 days of culture. Organoid models were only used to a maximum of passage 30. All cells were maintained in a humidified atmosphere containing 5% CO2 and tested negative for mycoplasma during monthly testing. The PPTO.46 organoid model used in this study was approved for use by the University of Toronto Research Ethics Board (Protocol no. 36107).

### Six-layer TRACER scaffold manufacturing

To seed ODCs, six-layer TRACER scaffolds first needed to be manufactured according to previously established protocols. Briefly, sheets of cellulose scaffold (Miniminit Products) were infiltrated with polymethyl methacrylate (PMMA) (Sigma-Aldrich), an oxygen impermeable polymer^35^. The PMMA-infiltrated scaffolds were then cut into individual strips (0.5 x 12 cm). Next, the seeding device was manufactured also following previously established protocols^35^. Briefly, PDMS Sylgard 184 (Dow Chemicals) was used to cast the base (20:1 w/w PDMS to crosslinker) and the lids (9:1 w/w PDMS to crosslinker) which were cured overnight at 80°C. Once cured, the inlet and outlet holes on the PDMS lids were created using a 1mm biopsy punch (Integra Life Sciences). Channel seeding guides were also created by cutting patterns in Parafilm™ (Bemis) sheets to accommodate for at least four six-layer scaffolds for seeding. The final devices were then assembled by placing the PDMS base on top of a polycarbonate backing (McMaster-Carr), layering the Parafilm™ channel seeding guides on top, and sandwiching TRACER scaffold strips between the base and the lids. Lids were properly aligned to the scaffold strips to allow for seeding. To sterilize the devices for seeding, each manufactured component was individually exposed to ultraviolet (UV) light for 15 minutes on each side, and then the assembled device was sterilized once more for another 15 minutes.

### Scaffold seeding for ODCs

To seed ODCs onto scaffolds, ODCs were first generated by dissociating Day 6-8 PPTO.46 PDOs with TrypLE (Gibco) dissociation solution for 30-40 minutes in an incubator. PDOs were mechanically dissociated with a micropipette every 10 minutes to ensure that single cells were generated. Once a cell suspension was obtained, ODCs were resuspended in a gel blend of 75:25 (v/v) neutralized Type I Bovine Collagen to Matrigel® at a cell density of 10 million cells/mL. To neutralize the Collagen, previously established protocols were used^32–34^. The gel blend and the cell density were previously determined by our group as the optimal parameters for seeding ODCs in TRACER^35^. The fully assembled and sterilized devices were then removed from the humidification chamber and placed on an ice pack to avoid premature gelation of the cell/gel mixture. To seed ODCs into the device, 5 µL of the ODC/gel suspension was injected into the inlet of each layer on the scaffolds (with the layer number side facing up). The ODC-infiltrated devices were then placed back into the humidification chamber and incubated for 45 minutes to allow for proper gelation. After the gelation period, the PDMS lids were removed, PBS was added on top of each strip to facilitate their detachment from the base, then each six-layer strip was transferred into a well-plate filled with media.

### Media optimization via CellTitre-Glo

To evaluate the impact of different media blends on ODC growth over time, 3000 ODCs were seeded in 10 µL domes of the gel blend (75:25 v/v neutralized Type I Bovine Collagen to Matrigel®) in an opaque, white 96-well plate compatible for luminescence readings. ODCs were grown in various media ratios of organoid CGM to RPMI-1640 (complete) for four days in an incubator; the ratios were 100:0, 75:25, 50:50, 25:75, 0:100 v/v. Four days was chosen because our group previously determined that this was the optimal timepoint for ODCs to form a mature, polarized, and confluent sheet in TRACER scaffolds. After four days, the media from ODCs domes were removed, PBS was added to wash away any residual media, and 100% RPMI-1640 (complete) was added to the domes and left to proliferate for another 4 days. As a control, ODCs grown in organoid CGM for 8 days were also included. To measure proliferation, the CellTiter-Glo 3D (Promega) assay was used according to manufacturer’s instructions. Briefly, every two days, an equal volume of CellTiter-Glo 3D was added to each well. To facilitate cell lysis, each dome was mechanically disrupted with a micropipette and left on a shaker platform for 30 minutes at room temperature and protected from light. Then, relative luminescence readings were obtained using a microplate reader. Luminescence readings were normalized to Day 0 readings. Since CellTitre-Glo 3D was an endpoint assay, multiple plates were seeded on the same day to measure luminescence every two days for eight days.

### Confocal Imaging of CK19 and ZO-1

To assess the expression of cytokeratin-19 (CK19) and zoncula occludens-1 (ZO-1) on ODCs cultured in different media blends, single layer ODC monocultures were seeded. Each single-layer ODC monoculture was cultured for four days in different media blends (100:0, 75:25, 50:50, 25:75, 0:100 v/v CGM to RPMI-complete), washed with PBS, and then cultured in 100% RPMI-complete for another two days. As a control, ODCs grown in organoid CGM for the whole experimental period were also included. After the six-day culture period, each sample was fixed with 4% paraformaldehyde (PFA) (Millipore Sigma) for 10 minutes at room temperature, then permeabilized with either 0.1% or 0.5% Triton X-100 (MilliporeSigma) for 5 minutes at room temperature for CK19 and ZO-1, respectively. Then, samples were blocked for one hour at room temperature with either 5% BSA with 0.2% Triton X-100 and 0.05% Tween-20 (MilliporeSigma) in PBS for anti-CK19 staining 5% BSA (MilliporeSigma) in PBS for anti-ZO-1 staining, as previously described^35^. For immunostaining, samples were incubated overnight at 4◦C with either anti-ZO1 (1:100; Clone ZO1-1A12, ThermoFisher Scientific) or anti-CK19 (1:400; Clone RCK108, Abcam) in their respective blocking solutions. The next day, secondary staining was performed with Alexa Fluor594 goat anti-mouse secondary antibody (1:400; ThermoFisher Scientific) for two hours at room temperature. Nuclei were counterstained with DRAQ5 (1:500, Cell Signalling Technology) in PBS for 10 min at room temperature. Then, scaffolds were mounted with Fluoromount-G (SouthernBiotech) and imaged with a Zeiss LSM 700 (Zeiss) confocal microscope equipped with a 20X objective (NA 0.8).

### Image analysis to measure CK19 expression

CK19 confocal images were imported and processed using the Fiji (National Institutes of Health) image analysis software. To obtain the mean gray value (MGV) of the CK19 signal, CK19 and DRAQ5 (nuclei) channels were first split, and a *Gaussian filter* (sigma = 1) was applied to smoothen the image and filter out extremely small dead cell fragments. Then, a mask was created to binarize the image (*Threshold, Max Entropy*) and nuclei were segmented using the *Watershed* function. Regions of interest (ROI) were then selected and the MGV of the CK19 signal surrounding each nucleus was obtained.

### Seeding TRACER co-cultures (single-layer and rolled)

On Day -6, CD14^+^ monocytes were magnetically sorted from freshly thawed PBMCS and differentiated to M0 macrophages for 6 days, using the methods described above. On Day -4, ODCs were seeded onto TRACER scaffolds (number-side facing up) using the methods for ODC seeding in TRACER and cultured in 75:25 v/v organoid CGM to RPMI-1640 (complete) for four days. On Day 0, ODC TRACER scaffolds were washed with PBS and placed number-side down on a PDMS seeding device, as previously manufactured^35^. Scaffolds were placed number-side down to facilitate contact between macrophages and ODCs. Differentiated M0 macrophages were then resuspended in RPMI-1640 (complete) media at a density of 80,000 macrophages/5 µL media. 5µL of macrophage cell suspension was then added directly on top of each layer of the ODC-containing scaffold strip (with number-side down) and placed in an incubator for 45 minutes. Then, each co-culture scaffold strip was transferred to a well-plate containing RPMI-1640 (complete) media for 24 hours. **For single layer strips**, co-cultures TRACERs were sectioned into individual layers and left in culture for one more day (total of 48 hours in co-culture) before downstream processing. **For rolled TRACERs,** scaffolds were carefully rolled around an aluminum mandrel and placed back into the well-plate containing RPMI-1640 (complete) media. The next day, co-culture TRACERs were rapidly unrolled for layer-specific analysis, with a total of 48-hours in co-culture and 24-hours in rolled co-culture.

### Seeding TRACER monocultures

**For rolled ODC monocultures,** six-layer TRACER scaffolds were seeded using the methods for ODC seeding in TRACER and cultured in 75:25 v/v organoid CGM to RPMI-1640 (complete) for four days. Then, ODC-containing scaffolds were washed once in PBS and transferred to a new well plate containing 100% RPMI-1640 (complete) at 37C. After 24 hours, the ODC-containing scaffolds were carefully rolled around an aluminum mandrel using the above method for rolling TRACERs. The next day, ODC-monoculture TRACERs were rapidly unrolled for layer-specific analysis. **For rolled macrophage monocultures,** M0 macrophages were differentiated from CD14+ PBMC-derived monocytes for 4-6 days as described above. M0 macrophages were then seeded onto TRACER scaffolds at a density of 16 million cells/mL (80,000 cells per 5 µL of gel blend) using the methods for ODC seeding in TRACER and incubated at 37C in RPMI-1640 (complete) media supplemented with a minimal amount of M-CSF (10 ng/mL) to ensure macrophage monocultures remained viable. After 24 hours, macrophage-containing scaffolds were rolled around an aluminum mandrel, placed in RPMI-1640 (complete) with 10 ng/mL M-CSF, and were subsequently unrolled for layer-specific analysis the next day.

### Confocal Imaging of fluorescently labeled TRACER co-cultures

To assess the localization of macrophages in co-culture with ODCs in TRACER scaffolds, single-layer GFP-PPTO.46 ODCs were seeded with CMTPX (Invitrogen)-labeled macrophages, using the described methods for seeding TRACER co-cultures. To label macrophages with CMTPX, M0 macrophages were incubated with 5µM of CMTPX in serum-free RPMI-1640 for 30 minutes at 37°C. Macrophages were pelleted by centrifugation, washed once, then resuspended in RPMI-1640 (complete) at a density of 80,000 macrophages/5 µL media for co-culture with ODCs. GFP PPTO.46 organoids were generated as previously described^35^. Single-layer fluorescently labeled cultures were then fixed with 4% PFA for 10 minutes at room temperature, mounted onto glass slides using Fluoromount-G, and imaged with a Zeiss LSM 700 confocal microscope equipped with a 20X objective (NA 0.8). Images were analyzed using Imaris software (Oxford Instruments).

### Digestion of co-cultures

To digest cells out of TRACER strips, an optimized digestion protocol was established. A new protocol was developed because we observed that the macrophages remained attached to the scaffold fibres and were not successfully digested out using the previously established protocols from our group. After evaluating various digestion cocktails (**Figure S9ai-iii**) with guidance from the Worthington Biochemical Corporation Manual^41,42^ and other studies^23^, we found that using a combination of Liberase TL (Roche) and Collagenase IV (Worthington Biochemical) was the most effective at digesting out ODCs and macrophages from the TRACER scaffolds based on both yield and viability. Briefly, each layer of TRACER (from single-layer or sectioned from rolled TRACER cultures) was washed with PBS to remove residual media and transferred to a 1.5 mL tube containing 280 µg/mL Liberase TL, 260 U/mL Collagenase IV and 260 U/mL DNAse I (MilliporeSigma). The tubes were placed on a Thermoblock at 37°C and 350 RPM for 45 minutes. Samples were immediately neutralized with media containing 10% FBS, then cells were pelleted by centrifugation at 300 G for 5 minutes, washed once with 5% FBS in PBS and centrifuged once more to obtain a cell pellet.

### EF5 analysis by flow cytometry

To measure oxygen levels in the different TRACER layers, flow cytometry of EF5 was performed as previously described^33,35^. Briefly, three hours prior to unrolling, 100 μM of EF5 (MilliporeSigma) was added to rolled TRACERs and incubated at 37°C. Then, TRACER constructs were unrolled, individual layers were sectioned, and digested using the protocol above. Digested cells were incubated with Zombie Violet (1:500) (BioLegend) to exclude dead cells from the analysis, fixed with 4% PFA, and blocked overnight at 4°C in PBS with 0.3% Tween-20, 5% normal mouse serum (MilliporeSigma) and 2% fat-free milk powder (BioShop). The next day, samples were stained with Cy3-conjugated anti-EF5 at 10 μg/mL for 3 h at room temperature. Flow cytometry was performed with a BD-Fortessa X20 flow cytometer. For controls, single-layer negative controls (without EF5 added), normoxic controls (not rolled, with EF5 added) and 0% pO2 anoxic controls (not rolled, exposed to an anoxic chamber overnight with EF5 added) were also included in the analysis. Anoxic controls were generated using an H85 Hypoxystation (Don Whitley Scientific) provided by the Wouters-Koritzinsky group at the University Health Network, Toronto, Canada. The median fluorescence intensity (MFI) of each sample was obtained and %EF5 binding was calculated as the ratio between the sample MFI subtracted by the negative control MFI, and the anoxic control MFI subtracted by the negative control MFI. The anoxic controls represented 100% EF5 binding.

### FACS sorting of co-cultures for gene expression analysis by qPCR

To analyze the gene expression of ODCs and macrophages, co-cultures were sorted via fluorescence activated cell sorting (FACS) with the assistance of the Princess Margaret Flow Cytometry Facility (University Health Network, Toronto). Macrophage-ODC TRACER co-cultures were digested using the protocol above to obtain a cell pellet from each sample. Then, samples were blocked with Human BD Fc Block™ solution (1 µL/200,000 cells) (BD Biosciences) for 10 minutes at room temperature to reduce any non-specific antibody staining by Fc Receptors expressed on macrophages. Samples were subsequently stained with Zombie Violet (1:400) viability dye and APC-Cy7 conjugated human anti-CD45 (BioLegend, clone: 2D1) for 45 minutes at 4°C protected from light. Viable Macrophages (Live^+^CD45^+^) and ODCs (Live^+^CD45^-^) were then sorted using a Beckman Mo-Flo Astrios cell sorter (Beckman Coulter) directly into 600 µL of RLT lysis buffer with 6 µL of β-mercaptoethanol. Samples were stored at -80°C if not immediately processed for RNA extraction.

To obtain RNA, samples were first homogenized using QIAshredder columns (Qiagen) and RNA was extracted using the Qiagen RNEasy Micro kit (Qiagen) according to the manufacturer’s instructions. RNA was quantified and reverse transcribed to cDNA using methods described above. 8 ng and 7 ng of macrophage and ODC cDNA was respectively used for qPCR, where gene-specific primers described in Table S2 were amplified and detected using the PerfeCTa SYBR® Green FastMix. The qPCR was performed using a CFX-384 thermocycler and data were acquired using Bio-Rad CFX Manager 3.1. Gene expression was normalized to the housekeeping genes 18S (for macrophages) and RPLP0 (for ODCs) and the 2^−ΔΔCt^ method was used to determine relative gene expression.

### Evaluating response to gemcitabine using alamarBlue Cell Viability assay

Gemcitabine-HCl (Sigma-Aldrich) was prepared in DMSO at a stock concentration of 10mM and single-use aliquots were generated and stored at -80°C and thawed as needed. To generate dose response curves, single layer ODC monocultures and ODC-macrophage co-cultures were manufactured as described above. Each co-culture layer was treated with varying doses (0, 20, 80, 140, 260, 300 µM) of gemcitabine for 72 hours. After 72 hours, each treated layer was incubated with 10% v/v alamarBlue solution in fresh RPMI-1640 (complete) media for 4 hours in an incubator. Then, 100 µL of each sample was transferred to a 96-well plate and fluorescence signal was analyzed using a standard fluorescence plate reader (αEx/Em = 560/590). The IC50 of ODC monocultures and co-cultures were calculated by fitting a 4-parameter logistics curve using GraphPad Prism 9 (GraphPad Software).

To investigate the impact of hypoxia on therapy response, rolled monoculture ODC and co-culture macrophage-ODC TRACERs were generated as described above. After 24 hours in rolled culture, TRACER constructs were either treated with 98 µM gemcitabine, the IC50 calculated for single-layer ODC monocultures, or with the vehicle control (0.98% DMSO) for 72 hours. Then, TRACERs were rapidly unrolled, individual layers were sectioned, and incubated with 10% v/v alamarBlue solution and analyzed using a fluorescence plate reader using the same method described above. Each layer was normalized to its corresponding DMSO-treated layer to obtain normalized cell viability for each layer (i.e., Layer 1 gemcitabine-treated co-culture was normalized to Layer 1 DMSO-treated co-culture). Normalized cell viability was then plotted against layer number as a surrogate for position, and a simple linear regression analysis was performed to determine whether the slope was significantly non-zero or significantly graded relative to a slope = 0.

### Collection of conditioned media from rolled TRACERs

Rolled ODC monocultures, macrophage monocultures and ODC-macrophage co-culture TRACERs were generated as described above. After 24 hours in rolled culture, TRACERs were rapidly unrolled and sectioned into individual layers. To collect conditioned media, layers one, three and six from each TRACER were placed in 600 µL of fresh RPMI-complete media. After 18 hours of incubation, conditioned media from each layer was collected into 1.5 mL tubes and centrifuged at 500 G for 5 minutes at 4°C to allow any cell debris to pellet at the bottom of each tube. After centrifugation, conditioned media was transferred to new 1.5 mL tubes by only taking media from the top portion and leaving around 50 µL of media to ensure that cell debris was not collected. Single-use aliquots of conditioned media were prepared and stored at -80°C until ready for use. As a positive control, single-layer macrophage monocultures, ODC monocultures, and ODC-macrophage co-cultures were incubated in fresh RPMI-1640 (complete) media for 18 hours in a 0.2% Hypoxia chamber (H45 Hypoxystation (Don Whitley Scientific) provided by the Wouters-Koritzinsky group at the University Health Network, Toronto, Canada).

### Evaluating response of T-cells to conditioned media (CM) collected from TRACER layers

To obtain fresh PBMC-derived T-cells, PBMCs were isolated from buffy coats as described above. T-cells were then obtained using a negative magnetic bead-based selection kit for pan T-cell isolation following the manufacturer’s protocol (Miltenyi Biotec). A negative selection was chosen to minimize any activation of T-cells during the selection process. To treat T-cells with conditioned media, T-cells were plated at a density of 300,000 cells per 300 µL of media (50:50 v/v conditioned media to fresh RPMI-complete) and incubated for 24 hours. The next day, plates were coated with 3µg/mL anti-CD3 (Invitrogen, clone: OKT3) for 2 hours at 37°C to prepare for T-cell activation. Then, T-cells that have been incubating with conditioned media were carefully transferred to the CD3-coated plates and further stimulated with 1 µg/mL anti-CD28 (BioLegend, clone: CD28.2) for another 48 hours. Five hours prior to the 48-hour timepoint, Brefeldin A solution (BioLegend) was added at a 1:1000 concentration to inhibit golgi-mediated secretion of proteins and to allow intracellular accumulation of cytokines. To optimize the assay, we also explored various protocols of T-cell activation based on methods previously described^43^, such as using PMA and Ionomycin to activate T-cells, and eventually proceeded with the method described here upon verifying that T-cell proliferation was observed after anti-CD3/CD28 activation (**Figures S9bi-ii**).

To analyze T-cells by flow cytometry, cells were collected and stained with Zombie Violet viability dye (1:400) for 10 minutes, then fixed and permeabilized with Foxp3 Transcription Factor Staining Buffer Set (eBioscience, Invitrogen) according to the manufacturer’s instructions. Samples were then stained for 30 minutes with: Human anti-CD3 (1:400; BioLegend, clone: SK7), human anti-CD4 (1:400; BioLegend, clone: OKT4), human anti-CD8 (1:400; BioLegend, clone: SK1), human anti-TNFα (1:400; BioLegend, clone: Mab11), human anti-IFNγ (1:400; BioLegend, clone: B27), human anti-GZMB (1:400; BioLegend, clone: QA18A28), human anti-Ki67 (1:200; BD Biosciences, clone: B56). Samples were analyzed by flow cytometry using a BD-Fortessa X20 flow cytometer.

### Statistical Analysis

Statistical analysis was performed using GraphPad Prism 9. p < 0.05 was considered significant. Power calculations were not employed to determine the number of replicates for each experiment. Details of the statistics used for each figure are summarized in the supplementary information (**Table S1**).

## 3. Results

### 3.1 Macrophages can retain plasticity to pro and anti-inflammatory phenotypes

Previous work in our group developed the TRACER2 platform, herein referred to as TRACER, which has been used to investigate the impact of hypoxic gradients on PDAC organoid-derived cell (ODC) monocultures^35^. In this study, we set out to adapt the TRACER system and incorporate primary macrophages for investigating macrophage-ODC interactions in hypoxic gradients. In developing these co-cultures, we first wanted to evaluate the feasibility of culturing primary macrophages in TRACER scaffolds, since primary macrophages have never been cultured in TRACER scaffolds and are highly plastic and sensitive cells whose phenotype is largely influenced by their local microenvironment. To determine whether the TRACER scaffold induced macrophage activation, we cultured unstimulated macrophages (“M0” macrophages) in 2D (tissue culture dish), gel blend domes (75:25 v/v% Type I bovine Collagen and Matrigel®), and single-layer TRACER scaffolds (TRACER scaffold + gel blend) (**Figure 1ai**), and then analyzed the expression of typical pro-inflammatory (IL-1β, IL-6, TNFα) and anti-inflammatory/pro-regenerative (CCL22, CD23, CD206) genes via qPCR^38–40^ We found that the relative mRNA expression of three out of the six genes (IL-6, CD23, CD206) were similar to 2D or gel alone cultures, while the other three genes, namely IL-1β (2D vs TRACER p-value= 0.0008; gel vs TRACER p-value = 0.0069), TNFα (2D vs TRACER p-value = 0.0045), and CCL22 (2D vs. TRACER p-value = 0.0018; gel vs. TRACER p-value = 0.0017) showed significant differences in expression compared to either 2D or gel alone cultures (**Figure 1aii**). Because there were differences in the relative expression of some genes, this suggested that there is likely a slight baseline activation of certain genes due to the TRACER scaffold fibres. Thus, it was important to next determine whether macrophages in the TRACER scaffold retained plasticity when activated with cytokines and whether this external stimulus overrode any baseline activation from the TRACER scaffold observed above.

**Figure 1:**
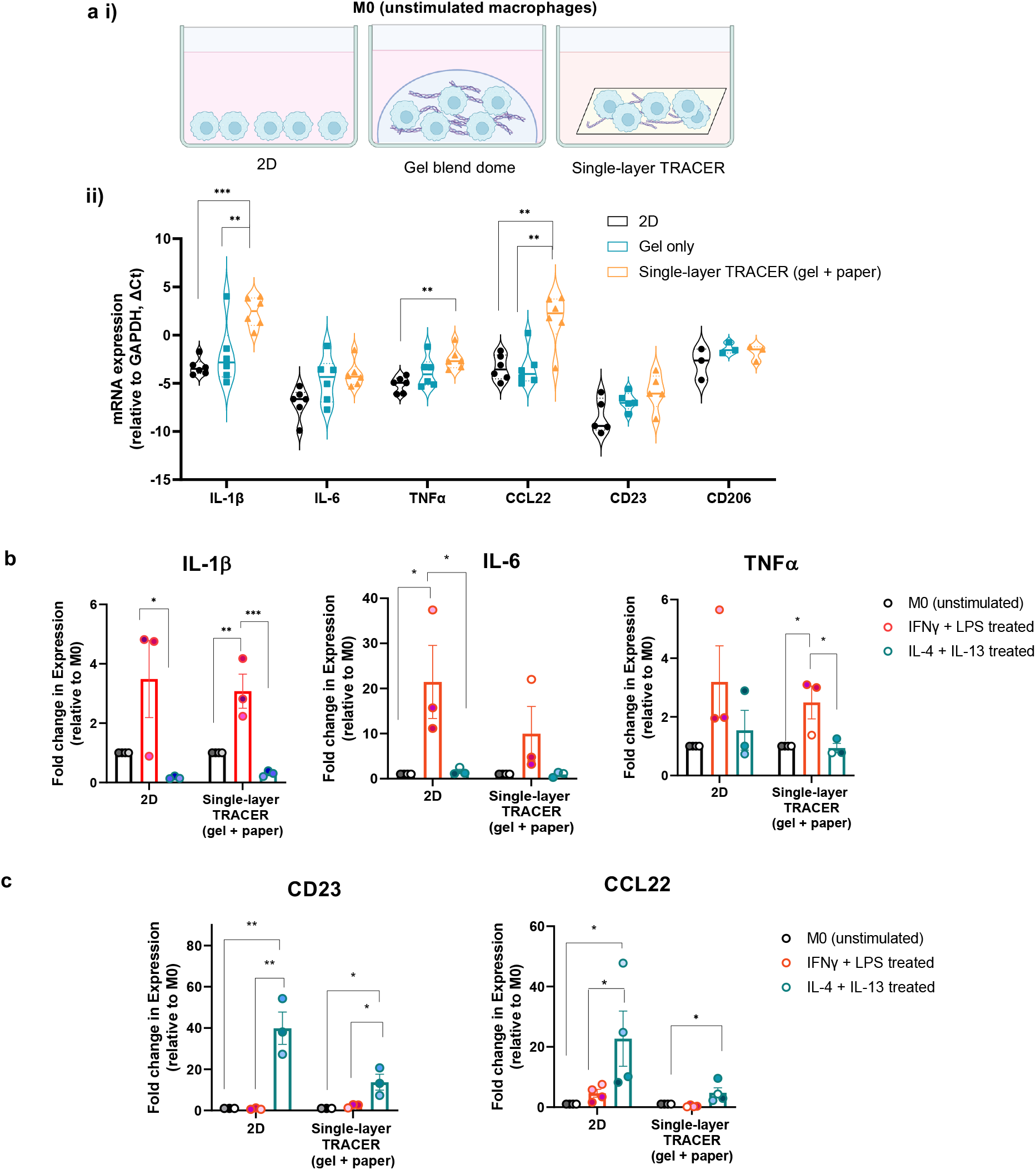
Primary macrophages are not significantly activated by the TRACER scaffold and retain plasticity to become M1 and M2 phenotypes when externally polarized. **(Ai)** Schematic of experimental set-up. Donor-derived macrophages were seeded on 2D, in a gel blend dome (75:25 v/v% Collagen to Matrigel ®) or infiltrated into single-layer TRACER scaffolds (gel blend and scaffold). Macrophages were left untreated (M0). **(Aii)** Quantitative PCR (qPCR) analysis of classic pro and anti-inflammatory genes in M0 macrophages cultured in 2D, gel blend dome, or single-layer TRACER scaffolds. Expression levels were quantified relative to the housekeeping gene GAPDH (βCt) (**p<0.01). **(B)** qPCR analysis of pro-inflammatory genes IL-6, IL-1β3, TNFα in unstimulated (M0), IFNγ/LPS and IL-4/IL-13 stimulated macrophages in 2D and single-layer TRACER scaffolds. Expression levels were quantified relative to the housekeeping gene GAPDH. Gene expression was normalized to the expression of M0 (unstimulated) macrophages in each condition and expressed as a fold change (ΔΔCt) (*p<0.05, **p<0.01, ***p<0.001). **(C)** qPCR analysis of anti-inflammatory genes CCL22 and CD23 in unstimulated (M0), IFNγ/LPS and IL-4/IL-13 stimulated macrophages in 2D and single-layer TRACER scaffolds. Expression levels were quantified relative to the housekeeping gene GAPDH. Gene expression was normalized to the expression of M0 (unstimulated) macrophages in each condition and expressed as a fold change (ΔΔCt) (*p<0.05, **p<0.01). Error bars are mean ± SEM for n= 3-6 independent donors (biological replicates).

Classically, the M1 (pro-inflammatory) and M2 (anti-inflammatory/pro-regenerative) dichotomy has been used to describe macrophage polarization states^38–40^. While this classification is now outdated, we used these simplified designations to investigate whether macrophages retained phenotypic plasticity in TRACER. Specifically, to evaluate macrophage plasticity in TRACER, we stimulated macrophages in 2D or cultured within a single TRACER layer with LPS and IFN-γ (M1; pro-inflammatory), IL-4 and IL-13 (M2; anti-inflammatory) or left them unstimulated (M0) and compared their responses. We also activated macrophages cultured in gel domes (**Figure S1a and S1b**), but found their gene expression to be highly variable, perhaps due to the heterogeneous distribution of cells in the gel dome versus the more evenly spread distribution in 2D and in TRACER scaffolds. Upon polarization, LPS and IFN-γ treated macrophages significantly upregulated the expression of pro-inflammatory genes IL-1β (M1 vs. M2 p-value **=** 0.028) and IL-6 (M1 vs. M2 p-value **=** 0.022; M1 vs. M0 p-value = 0.034) in the 2D configuration, and IL-1β (M1 vs M2 p-value = 0.001; M1 vs. M0 p-value = 0.0036) and TNFα (M1 vs M2 p-value = 0.0226;M1 vs M0 p-value = 0.02) in the TRACER scaffold (single TRACER layer) (**Figure 1b**). Similarly, IL-4 and IL-13 treated macrophages significantly upregulated the expression of anti-inflammatory genes CCL22 and CD23 in the 2D (M2 vs M0 p-value = 0.0432; M1 vs. M2 p-value = 0.091 for CCL22; M2 vs. M0 p-value = 0.022 and M2 vs. M1 p-value = 0.021 for CD23) and single-layer TRACER scaffolds (M1 vs M2 p-value = 0.0254 for CCL22; M2 vs M0 p-value = 0.0167 and M2 vs M1 p-value = 0.0262 for CD23) (**Figure 1c**). Although we observed some variability in the results likely due to donor variation, macrophages showed the expected expression of pro and anti-inflammatory genes upon activation in single TRACER layer strips. Thus, we concluded that the macrophages retained plasticity in single-layer TRACER scaffolds and that any baseline effect of the TRACER scaffold (as observed for IL-1β, TNFα and CCL22) was minimal relative to gene expression changes induced by external activation. Overall, our results suggested that TRACER is a feasible platform to evaluate macrophage phenotypic activation and that any changes driven by external stimulation, such as cytokine gradients or eventual co-culture with tumor cells, can be observed in TRACER.

### 3.2 Development of a co-culture media regime to ensure that ODCs maintain proliferation, maturity, and polarization for eventual co-culture with macrophages

In the development of TRACER, we previously found that growing ODCs at a density of 10M cells/mL in a gel blend of 75:25 v/v% Type I bovine Collagen and Matrigel® for four days in complete organoid growth media (CGM) was sufficient for ODCs to mature, polarize and form a confluent sheet in the TRACER scaffolds^35^. Therefore, we set day four as the timepoint for incorporating macrophages, which mimicked the *in vivo* situation wherein macrophages are recruited to the TME by mature and well-differentiated tumors^44^. However, before we could incorporate macrophages with ODCs to generate TRACER co-cultures, we set out to determine the optimal co-culture media to enable both cell types to function appropriately. Specifically, PDOs are typically grown in organoid CGM, a defined media composed of several small molecules (TGFβ inhibitor, Rho-kinase inhibitor, nicotinamide) and mitogens (FGF-10, EGF, Noggin) required for organoid initiation and maintenance^45–49^. On the other hand, PBMC-derived macrophages are maintained in a basal media of RPMI-1640 (complete). We speculated that culturing ODCs in 100% RPMI-1640 (complete) could negatively impact their growth and ability to form organoids due to the lack of necessary growth factors and therefore eliminated the idea of performing all experiments in RPMI-1640 alone. We also speculated that culturing macrophages in 100% organoid media could unfavorably activate these highly plastic myeloid cells. As expected, upon culturing PBMC-derived macrophages in organoid media, cells displayed a mixed pro- and anti-inflammatory phenotype, suggesting that small molecules and mitogenic factors in the organoid media unfavorably activated the macrophages (**Figure S2a**).

Given that premature activation of macrophages is not ideal and could impact or limit the detection of tumor cell influences on macrophage phenotype when the two cell types are in co-culture, we sought to develop a co-culture media composition which supported ODCs growth and maturation but did not result in macrophages being prematurely activated by the media. As a strategy to achieve this, we investigated whether pre-conditioning ODCs in a media blend before co-culturing them with macrophages in a growth-factor free media of RPMI-1640 (complete) would be a viable approach. To do this, we cultured gel domes of ODCs in varying blends of organoid CGM and RPMI-1640 (complete) (100:0, 75:25, 50:50, 25:75, 0:100 v/v%) media for four days, the time needed for organoids to grow and mature as established previously^35^. Then, we removed and washed away any remaining media, and replaced with fresh 100% RPMI-1640 (complete) media for an additional four days. Using the CellTiter-Glo 3D proliferation assay, we found that pre-conditioning ODCs in varying media blends before switching to 100% RPMI-1640 (complete) all resulted in similar growth dynamics compared to culturing ODCs in its standard media (100% organoid CGM) for the whole assay duration (**Figure 2a**). This implied that growing ODCs in a media blend for four days before totally switching to RPMI-1640 (complete) was a potentially viable strategy to maintain ODC growth while transitioning the cells into a media compatible with macrophage culture. Culturing ODCs in 100% RPMI-1640 (complete) immediately after TRACER seeding however resulted in significantly slower growth dynamics compared to all the other conditions (p-value = 0.0039), suggesting that ODCs need some component present in the organoid media to initiate and maintain growth (**Figure 2a**).

**Figure 2:**
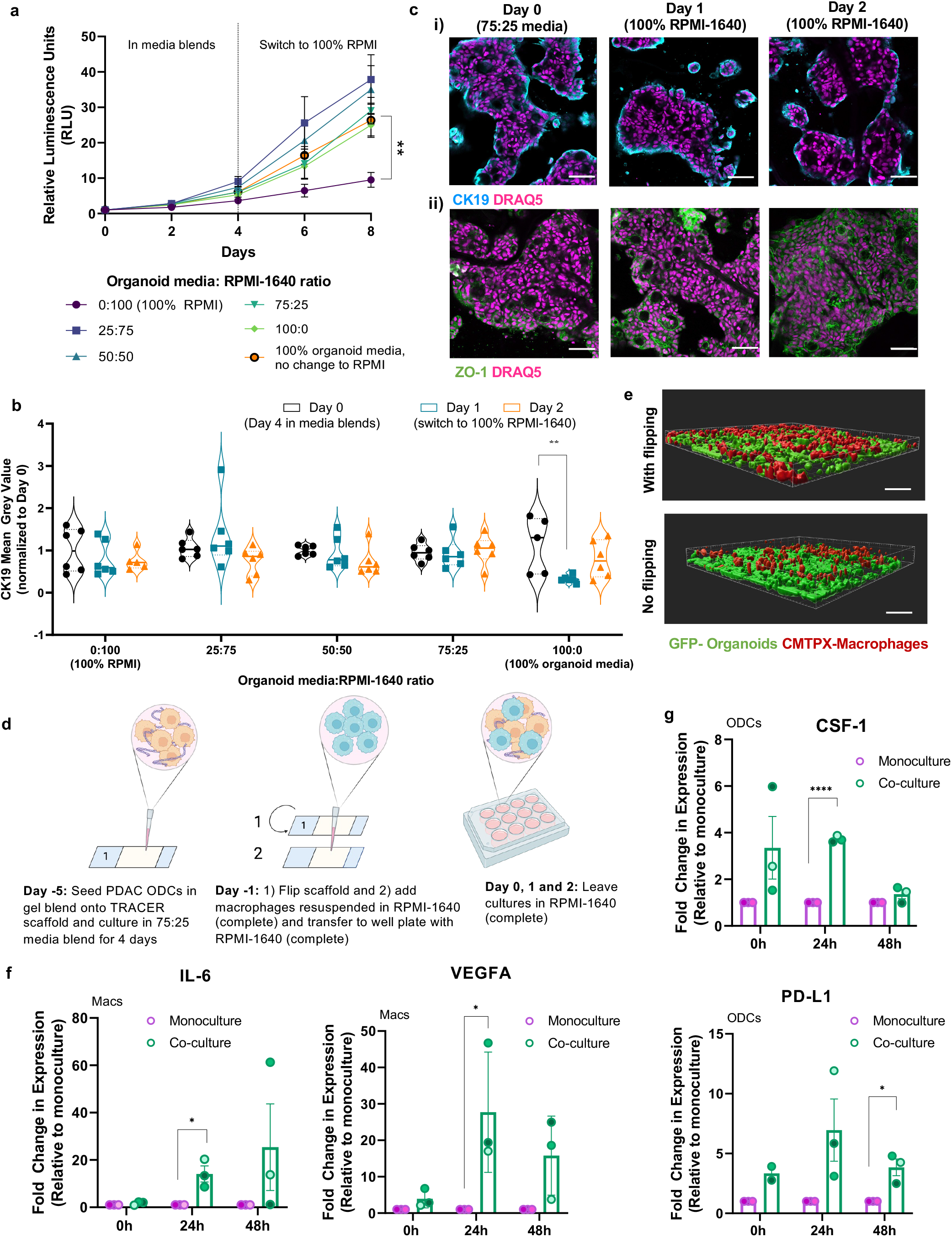
Development of an optimized primary macrophage and ODC co-culture strategy in non-rolled TRACER co-culture. **(A)** Proliferation of ODCs over time measured using the luminescence-based CellTiter-Glo 3D assay every 2 days for a total of 8 days. The assay read-out was displayed as relative luminescence (RLU) over time (p**<0.01). **(B)** Quantification of CK19 expression (mean grey value (MGV)) of ODCs in single-layer scaffolds, measured by immunofluorescence and confocal microscopy at Days 0, 1 and 2. ODCs were pre-conditioned in varying media blends for 4 days (Day 0) before replacing the media with 100% RPMI-1640 (complete) in all conditions, except for the control group (no change to RPMI). CK19 expression was normalized to Day 0 (p**<0.01). n=3 biological replicates with 2-3 technical replicates each. **(C)** Representative confocal images of CK19 and ZO-1 expression of ODCs in single-layer TRACER scaffolds at Day 0, 1 and 2. Scale bar = 100µm. **(D)** Schematic of macrophage-ODC co-culture strategy in single-layer TRACER scaffolds. Day 0, 1 and 2 represents 0h, 24h and 48h in Figures 2F and 2G. **(E)** Confocal images of GFP-ODCs co-cultured for 24h with CMTPX-labeled macrophages added onto the ODC-containing scaffolds with or without flipping. Scale bar = 100µm. **(F)** qPCR of pro- and anti-inflammatory genes, IL-6 and VEGFA respectively, of macrophages in co-culture with ODCs versus monoculture macrophages at 0h, 24h and 48h. The 0h timepoint represents the 24h after macrophages were added onto the ODC-containing scaffolds for co-culture. Gene expression was normalized to the housekeeping gene 18S and expressed as a fold change relative to monoculture macrophage controls (p*<0.05). **(G)** qPCR of CSF-1 and PD-L1 expression of ODCs in co-culture with macrophages versus monoculture ODCs at 0, 24 and 48h. The 0h timepoint represents the 24h after macrophages were added onto the ODC-containing scaffolds for co-culture. Gene expression was normalized to the housekeeping gene RPLP0 and expressed as a fold change relative to monoculture ODC controls (p*<0.05, p****<0.0001). Error bars are mean ± SEM for n= 3 biological replicates (for ODCs) or n= 3 independent donors (for macrophages).

To further guide our selection of which media blend to use for ODC pre-conditioning, we visualized the expression of Cytokeratin-19 (CK19), a PDAC-specific epithelial marker^20,45,50^, in single-layer TRACER ODCs via confocal imaging. We observed that CK19 expression was maintained for two days after switching to 100% RPMI-1640 (complete) for all media blend compositions, except for the 100% organoid media condition where we noted a significant drop in CK19 levels (p-value = 0.0026) one day after the media switch (**Figure 2b**, **representative images in Figure S2b**). Notably, we observed the most stable CK19 expression over two days in the 75:25 v/v% blend (**Figure 2b and 2ci**). For this specific 75:25 v/v% media blend strategy, we also visualized the expression of zoncula occludens-1 (ZO-1) in single-layer ODC TRACERs; ZO-1 is a tight-junction marker whose localization acts as an indicator of polarity and lumen formation^49,51,52^. We found that ZO-1 localized apicobasally in ODCs pre-conditioned with the 75:25 media blend, indicating lumen formation occurred both during the four-day initial culturing period and after switching to 100% RPMI-1640 (complete) (**Figure 2cii, representative images in Figure S2c**). Together with the stable expression of CK19, these data suggested that conditioning ODCs in the 75:25 v/v% media blend was a viable strategy for maintaining organoid maturation and polarization for at least two days after replacing the media to RPMI-1640 (complete).

Recent studies have reported that PDAC organoids phenotypes are sensitive to media composition and exhibit a phenotype resembling the aggressive basal-like PDAC disease when cultured in either RPMI-1640 (complete) or organoid media stripped of certain growth components; in contrast, organoids cultured in CGM resembled the classical PDAC subtype^53^. Since our selected media strategy involved transitioning organoids into culturing in RPMI-1640, we assessed the impact of the media switch on the expression of classical and basal markers in TRACER. Consistent with this study^53^, we observed that ODCs upregulated basal-disease markers (KRT17 and WNT7B) when cultured in either RPMI-1640 (complete) or the 75:25 v/v% media blend and downregulated classical-disease markers (LYZ and GATA6) although only when cultured in RPMI-1640 (complete) (**Figure S2di and S2dii**). Further consistent with this idea, macrophages downregulated the TAM marker SPP1 (not statistically significant) which is associated with classical PDAC, when exposed to conditioned-media collected from ODCs grown in either RPMI-1640 (complete) or the 75:25 blend (**Figure S2e**). Together, these findings are consistent with the Raghavan *et al* study and suggest that the use of our media blending and switching strategy likely models more aggressive and chemo-resistant basal-like PDAC more closely than the classical subtype^53^. More extensive genomic and transcriptomic analysis is needed, however, to verify this.

In summary, our data indicate that pre-conditioning ODCs for four days before switching to RPMI-1640 (complete) with a 75:25 media blend was a viable strategy for subsequent co-culture of ODCs with macrophages in TRACER. The pre-conditioning strategy maximized the initial ratio of CGM that ODCs were exposed to, allowed ODCs to maintain growth, maturation, polarity, and potentially models the more aggressive basal-like PDAC subtype.

### 3.3 Optimization of a direct co-culture method for macrophages and ODCs in TRACER scaffolds reveals that 24-hours in co-culture is sufficient to induce phenotypic changes

Having determined the optimal co-culture media and media conditioning regime, we next set out to optimize a method for directly co-culturing macrophages and ODCs in TRACER. To do this, we first incorporated primary macrophages into TRACER scaffolds infiltrated with ODCs that had been matured for 4 days by directly adding donor-derived macrophages suspended in RPMI-1640 (complete) onto single-layer TRACER scaffolds (**Figure 2d**). The number of macrophages that were added to each layer represented around 10-20% of the number of ODCs after 4 days of culture (**Figure S3a**), mimicking the estimated *in vivo* proportions of macrophages in PDAC^5,54^. We then incubated the single-layer ODC-macrophage scaffolds for 45 minutes before transferring each single-layer scaffold to RPMI-1640 (complete) media (**Figure 2d**). To allow the macrophages to sufficiently infiltrate, we found that flipping the paper scaffold (number-side down) before directly co-culturing the primary macrophages resulted in better cell-cell contact between the ODCs and macrophages after 24 hours (**Figures 2d and 2e**). The scaffold flipping strategy was previously adopted by our group^55^ to incorporate a second cell type into a scaffold and is likely because the ODCs are observed to settle to one side of the cellulose scaffold during cell-gel infiltration (data not shown). Interestingly, after 48 hours, the ODCs and macrophages were observed to localize in the same z-plane in both “flipped” and “non-flipped” conditions (although infiltration was still more obvious in the “flipped” condition), indicating that macrophages could sufficiently infiltrate the scaffold over a 48-hour timeframe when in co-culture (**Figure S3b**). However, to simplify the protocol and minimize the assay duration, we chose to flip the scaffolds and designate 24 hours as the minimum time needed for macrophages and ODCs to spatially encounter each other.

Since TRACER is best suited to endpoint read-outs, we next set out to determine a reasonable interaction time necessary to produce phenotypic changes in the ODC and macrophages cell populations, which would enable us to select an appropriate endpoint time to unroll the TRACER co-cultures. To do this, we co-cultured macrophages and ODCs together for 0, 24 and 48 hours in single-layer TRACER scaffolds, with 0 hours representing the 24 hours after the scaffold is flipped to allow for sufficient macrophage-ODC contact. At each time point, co-cultures were digested using an optimized cocktail of Liberase TL and Collagenase IV (**Figure S9ai-iii**) then FACS-sorted via CD45^+^ (macrophages) and CD45^-^ (ODCs) directly into lysis buffer for subsequent RNA extraction and qPCR analysis. First, we considered the impact of the ODCs on the macrophage cells and found that macrophages displayed >20-fold increases in expression of VEGFA (p value = 0.049) and IL-6 (p-value = 0.0181) as early as 24 hours in co-culture compared to monoculture (**Figure 2f)**. These expression increases after 24h of co-culture are consistent with induction of a TAM-like phenotype as IL-6 and VEGFA are respectively associated with inflammatory and angiogenic TAMs^56–58;^ expression changes in other TAM markers at the 24h timepoint were more variable. For example, CCL22 and IL-1β were more prominently upregulated (not statistically significant) at 24 and 48 hours respectively, while little to no change in CD206 expression was observed across all timepoints in macrophage co-cultures versus monocultures (**Figure S3c**). The variable expression of these genes highlighted that evaluating TAM phenotype with only a few markers is challenging, that there is significant donor variation, and that gene expression is potentially cyclic and fluctuates over time. While fully interpreting macrophage phenotypes will therefore require more granular analysis such as transcriptomic or proteomic strategies, our data indicated that ODCs induced phenotypic changes in macrophages after 24-hours of co-culture, and this is a reasonable timepoint to assess rolled TRACER co-cultures.

We next considered the response of ODCs in response to co-culture with macrophages. We observed that ODCs showed an approximately 4-fold increase in the expression of colony stimulating factor-1 (CSF-1), a critical macrophage survival and differentiation factor^59^, at 0 hours (i.e., after 24h of co-culture to allow the macrophages to integrate into the tissue) (not statistically significant) and 24 hours (p-value <0.0001) compared to monoculture ODCs (**Figure 2g**). Increased expression of CSF-1 as early as 0 hours suggested that macrophages in co-culture may promote their own survival via CSF-1 through rapid, likely soluble factor interactions with ODCs via either direct or paracrine signals. This observation is consistent with previous studies which have shown that TAMs can induce CSF-1 expression in cancer cells to promote their own differentiation and survival^59,60^. Interestingly, increases in CSF-1 expression in co-cultures were lost by the 48-hour timepoint, supporting the notion that gene expression is cyclic and varies over time. In addition to CSF-1, the expression of programmed death ligand-1 (PD-L1), a checkpoint inhibitor to help T-cells distinguish self from non-self, increased in ODCs after 0, 24 and 48 hours in co-culture when compared to ODCs in monoculture, although statistical significance was only observed at the 48-hour timepoint (p-value = 0.015). This observation was consistent with previous reports in which macrophages induced the upregulation of PD-L1 in tumours, which has been correlated with an increased immunosuppressive phenotype in various cancers^61–63^. Based on these observations of both ODCs and macrophages we determined that 24 hours after the initial cell-contact period in single-layer TRACERs appeared to be sufficient to induce observable phenotypic changes in either cell type. We therefore selected a timepoint of 24h after TRACER rolling to assess the impact of co-cultures for all future experiments.

### 3.4 Rolled macrophage-ODC TRACER co-cultures generate hypoxic gradients and exhibit expected graded responses to hypoxia

Having determined the media strategy and duration of rolled co-culture, we defined the final workflow for co-culturing macrophages and ODCs in TRACER (**Figure 3a**). Briefly, PDAC ODCs were seeded onto six-layer TRACER scaffolds and cultured in a media blend of 75:25 v/v% organoid CGM and RPMI-1640 (complete) for 4 days. Then, the scaffold was flipped, macrophages were added, and left unrolled for 24 hours in 100% RPMI-1640 (complete) to allow for cell-cell contact. After 24 hours, TRACER co-cultures were rolled around an oxygen impermeable TRACER mandrel for another 24 hours before rapidly unrolling for layer specific analysis. To distinguish macrophages from ODCs during analysis, macrophages were either labelled with a CellTracker dye (CMTPX) or with a fluorescently tagged antibody against the pan-immune cell marker, CD45.

**Figure 3.**
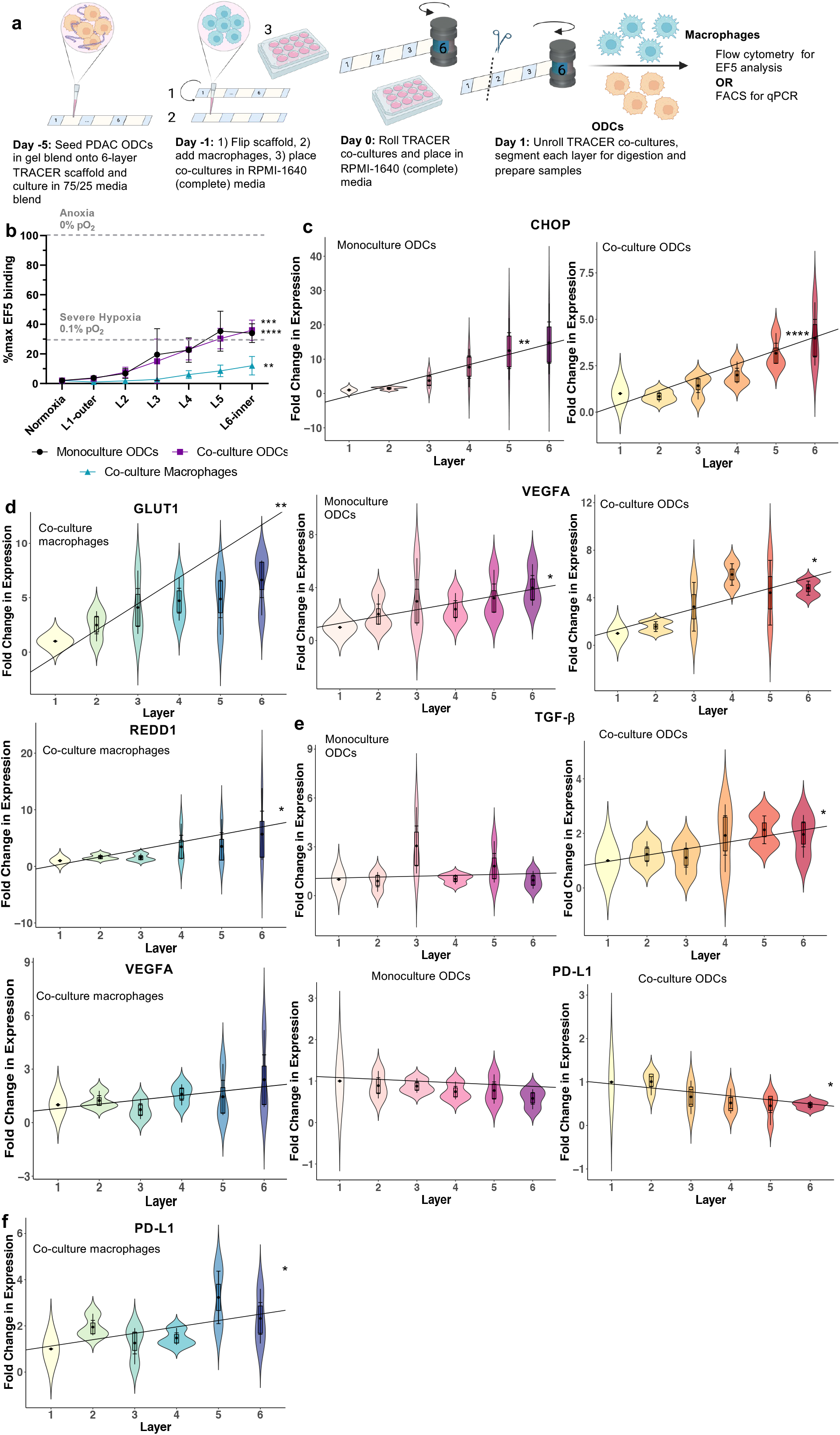
Measuring graded hypoxia response in rolled macrophage-ODC TRACER co-cultures. **(A)** Schematic of rolled macrophage-ODC co-culture in TRACER. **(B)** Quantification of EF5 binding 24h after rolling of ODC monocultures, ODCs in co-culture with macrophages and macrophages in co-culture with ODCs revealed the presence of oxygen gradients across the TRACER layers. EF5 binding is displayed as a percentage of the maximum EF5 binding (relative to anoxia controls). Grey lines correspond to theoretical EF5 binding for a given oxygen concentration^64,67^. (EF5 binding vs Layer 1 (outer layer): p**<0.01, p***< 0.001, p****<0.0001). Error bars are mean ± SEM for n= 3-4 biological replicates. **(C)** qPCR of hypoxia response genes CHOP and VEGFA in 24h rolled ODC co-cultures versus ODC monocultures. Gene expression levels were relative to the housekeeping gene RPLP0 and expressed as a fold change normalized to Layer 1 (outer layer). (slope of best fit line vs. 0: p*<0.05, p**<0.01, p***< 0.001). **(D)** qPCR of immune regulatory genes TGFb and PDL1 in 24h rolled ODC co-cultures versus ODC monocultures. Gene expression levels were relative to the housekeeping gene RPLP0 and expressed as a fold change normalized to Layer 1 (outer layer). (slope of best fit line vs. 0: p*<0.05, p**<0.01, p***< 0.001). **(E)** qPCR of hypoxia response genes VEGFA and REDD1 in 24h rolled macrophage co-cultures. Gene expression levels were relative to the housekeeping gene 18S and expressed as a fold change normalized to Layer 1 (outer layer). (slope of best fit line vs. 0: p*<0.05, p**<0.01, p***< 0.001). **(F)** qPCR of immune regulatory gene PD-L1 in 24h rolled macrophage co-cultures. Gene expression levels were relative to the housekeeping gene 18S and expressed as a fold change normalized to Layer 1 (outer layer). (slope of best fit line vs. 0: p*<0.05, p**<0.01, p***< 0.001). Error bars of boxplots in violin plots for are mean ± SEM for n= 3-4 biological replicates (for ODCs) or n= 3 independent donors (for macrophages).

With the complete co-culture protocol established, we set out to investigate whether hypoxic gradients existed in rolled TRACER co-cultures. To measure hypoxic gradients, we added EF5, a small molecule that is metabolized and forms covalently bound adducts in hypoxic cells^64–67^, to the rolled cultures 3 hours prior to unrolling. After unrolling, we digested cells from each layer and stained with an anti-EF5 antibody for analysis via flow cytometry. To distinguish the impact of macrophages versus the gradient generated, rolled ODC monoculture TRACERs were included as a point of comparison. As expected, we observed hypoxic gradients in ODCs in monoculture and in co-culture with macrophages (slope of regression line different from 0: p = 0.0007 for monoculture; p = 0.0001 for co-culture) from layer one to six in TRACER, with an increase in EF5 binding from 0% (normoxia) to around 30% (severe hypoxia, 0.1% pO2) (**Figure 3b**). Unsurprisingly, the gradients generated by ODCs in monoculture and co-culture had a similar gradient and apex point (30% EF5 binding), suggesting that macrophages did not greatly impact the oxygen gradient. This suggested that, in comparison to macrophages, ODCs were the dominant drivers that established the oxygen gradient, which was consistent with the fact that cancer cells are more metabolically active and have higher oxygen consumption rates than macrophages^68–70^. When we measured EF5 levels in TRACERs generated from macrophages alone we did not observe a hypoxic gradient (**Figure S4a**). However, macrophages did experience hypoxia when in co-culture with ODCs in TRACER, indicating that hypoxic reprogramming of macrophages was present in co-culture TRACERs (**Figure 3b**). Specifically, when we measured the hypoxia levels of macrophages co-cultured with ODCs in TRACER, we observed a significantly increasing hypoxic gradient from layer one to six (slope of regression line different from 0: p **=** 0.007). However, EF5 binding only increased from 0% to 10% (moderate hypoxia, 0.5% pO2) which was a less steep gradient than observed in ODCs from co-cultures (**Figure 3b**) and can likely be attributed to the larger oxygen consumption rate of tumour cells compared to macrophages^68–70^. Together, our observations suggested that hypoxic gradients driven by ODC consumption were present in TRACER co-cultures.

We next verified that ODCs and macrophages in TRACER displayed expected transcriptional responses to hypoxic gradients by performing qPCR on FACS sorted ODCs (negative anti-CD45 signal) and macrophages (positive anti-CD45 signal). In concordance with ODCs in monoculture, ODCs co-cultured with macrophages exhibited a statistically significant increasing graded expression in C/EBP Homologous Protein (CHOP) (slope of regression line different from 0: p-value **=** 0.0011 for monoculture; p-value <0.0001 for co-culture) and vascular endothelial growth factor alpha (VEGFA) (slope of regression line different from 0: p-value **=** 0.0254 for monoculture; p-value = 0.0238 for co-culture), which are both known hypoxia response genes involved in the unfolded protein response^71,72^ and angiogenesis^73,74^, respectively (**Figure 3c**). These observations were consistent with previous TRACER studies that evaluated transcriptional response of CHOP and VEGFA in the same ODC line (PPTO.46) used in this study^35^ in monoculture. Similarly, we observed expected gradients in the expression of hypoxia response genes in macrophages in co-culture with ODCs. Specifically we observed gradients in glucose transporter protein type 1 (GLUT1) (slope of regression line different from 0: p value **=** 0.0047), regulated in development and DNA damage responses 1 (REDD1) (slope of regression line different from 0: p value **=** 0.0469) and VEGFA (not significant; slope of regression line different from 0: p value **=** 0.862), known hypoxia response genes involved in glycolysis^75,76^, DNA damage^77,78^, and an angiogenic TAM-like phenotypes^58,79,80^, respectively (**Figure 3d**). Note that unlike ODCs, we did not compare the expression profile of macrophage TRACER co-cultures with their corresponding TRACER monocultures because macrophage monocultures did not generate hypoxic gradients in TRACERs (**Figure S4a).** Importantly, these observations that hypoxia response genes were expressed in both ODCs and macrophages suggested that the hypoxic gradients generated in TRACER co-cultures were sufficient to induce a known graded transcriptional change to hypoxia and that the cells in the co-culture TRACER were behaving as expected.

### 3.5 Hypoxic reprogramming of macrophage and ODC phenotypes is evident in TRACER

Having established that macrophages and ODCs showed the expected transcriptional changes to cell-derived oxygen gradients in TRACER, we next wanted to investigate more extensively the impact of hypoxia on macrophage and ODC gene expression. In ODCs, we measured the expression of the immune-regulatory genes transforming growth factor beta (TGFβ) and PD-L1. TGFβ in PDAC has been shown to facilitate EMT, induce immunosuppressive T-cells, myeloid cells, and fibroblasts, and overall correlate with poor prognosis and the basal PDAC subtype^81,82^. In monoculture TRACERs, ODCs exhibited a slight increase in TGFβ expression in layer three (moderate hypoxia), but no overall change in expression from layers one to six (**Figure 3e**). However, in co-culture with macrophages, ODCs displayed a shift from the flat profile observed in monoculture to increasing graded expression of TGFβ in the ODC from co-cultures (slope of regression line different from 0: p-value = 0.0279) (**Figure 3e**). This increased expression of TGFβ by ODCs in hypoxic co-culture is consistent with studies showing that TGFβ secretion is increased in hypoxic pancreatic cancer cells^83,84^. We also measured expression of PD-L1 in TRACER co-cultures. PD-L1 is a ligand that binds to PD-1 on T-cells, is a checkpoint marker for discriminating self from non-self and has been associated with immune evasion in many cancers including PDAC^85^. Intriguingly, we found that PD-L1 expression in rolled monoculture ODCs remained consistent from layer one to six but decreased in the inner layers of rolled co-culture ODCs (slope of regression line different from 0: p-value **=** 0.0267) (**Figure 3e**). Similar to our observations in TGFβ expression, we noted a shift from a flat expression profile of PD-L1 in monoculture to a decreasingly graded expression profile in co-culture. This further supported the notion that hypoxic macrophages resulted in reprogramming of ODC phenotype in TRACER co-cultures over the timeframe of our experiments. We note that our observation of decreased PD-L1 expression in the inner layers of the TRACER co-culture conflict with previous studies which reported that PD-L1 levels were elevated in hypoxic cancer cells^86–88^. This inconsistency is potentially due to different cellular responses in the controlled hypoxic chambers used in these studies versus in TRACER, wherein ODCs were additionally exposed to cell-generated gradients of other molecules (nutrients and waste) in addition to hypoxia. Interestingly, we found that the fold-change in CHOP and TGFβ expression was elevated in ODCs retrieved from layer six of the TRACER co-culture (relative to layer one) compared to ODC co-cultures placed in a 0.2% hypoxia chamber (relative to normoxia controls) (**Figure S4b**). Alternatively, when we compared the fold change in expression for VEGFA and PD-L1, unlike CHOP and TGFβ expression, we observed similar results wherein PD-L1 and VEGFA expression respectively decreased and increased in both layer six TRACERs and the hypoxic controls (**Figure S4b**). Our data therefore suggest that at least some expression patterns, such as CHOP and TGFβ observed here, are uniquely present in TRACER, and are potentially dominated by cell-derived gradients, such as metabolic and molecular gradients, rather than hypoxia alone. This observation highlights a potential value of the TRACER culture platform over simply culturing cells in a controlled hypoxia chamber.

Notably, the absence of graded PD-L1 expression in monoculture ODCs also conflicted with our previous work in TRACER in which we observed increases in PD-L1 expression in rolled monoculture ODCs layer one to six^35^. These conflicting results however are likely explained by differences in culture media used between the two studies. In our previous work, ODC monoculture TRACERs were cultured in complete organoid growth media for the duration of the study, while in this study, ODC monoculture TRACERs were cultured in a media blend first before switching to 100% RPMI-1640 (complete). The media switch effectively removed organoid growth components to render the co-culture more compatible with macrophages but resulted in an increased expression of some basal-like genes (**Figure S2d**) and an elevated baseline expression of PD-L1, as well as other genes, in normoxic cultures (**Figure S5a**). These data suggest that the influence of hypoxic gradients on basal disease potentially may not be as prominent compared to the impact of hypoxia on classical disease and that media differences in PDAC ODC culture are significant in dictating response to graded hypoxia.

We next evaluated macrophage gene expression patterns in TRACER co-cultures and observed increased expression of the checkpoint inhibitor PD-L1 from layer one to six (slope of regression line different from 0: p-value **=** 0.0436) (**Figure 3f**) in the macrophages. This was consistent with previous studies which reported that murine macrophages upregulated PD-L1 when placed in a hypoxia chamber^89^ and that PDAC patient TAMs displayed elevated levels of PD-L1 at the protein level^56^, although a link between PD-L1 and hypoxia was not investigated in the second study. While these two studies are consistent with the observations in **Figure 3f**, the increased expression of PD-L1 in our TRACER co-culture system is the first to explicitly suggest a connection between elevated PD-L1, cancer cell-generated hypoxic gradients, and primary macrophages; this highlights the value of TRACER in potentially exploring more detailed mechanisms of cancer-immune cell interactions in hypoxic gradients. We also quantified the expression of IL-1β and IL-6 in co-culture TRACER macrophages, and expected to see an increased expression of these two genes based on previous studies^90–92^. However, we did not observe any significant graded change in any of these genes in TRACER (**Figure S4c**) perhaps due to donor variation and the inherent difficulty of interpreting heterogenous macrophage phenotypes using qPCR. Despite these variations, our data suggested that the expression of some macrophage and ODC genes are uniquely rewired by cell-generated hypoxic gradients in TRACER, highlighting the value of the TRACER system in deciphering hypoxia reprogrammed interactions between macrophages and cancer cells.

### 3.6 Evaluating response to therapy in TRACER co-cultures reveal that macrophages chemo-protect ODCs from gemcitabine in hypoxia

TAMs in PDAC have been shown to contribute to resistance of the standard-of-care PDAC chemotherapeutic drug, gemcitabine. Specifically, tumour-educated macrophages secrete deoxycytidine, a gemcitabine analog that competitively inhibits gemcitabine uptake^93^, and produce deoxycytidine kinase which drives cancer cells to metabolize gemcitabine to its inactive form^94^. While there have been many studies investigating the mechanisms of TAM-mediated gemcitabine resistance in normoxia, to our knowledge, none have explored TAM-mediated gemcitabine resistance in graded hypoxia. Thus, to investigate whether hypoxia plays a role in driving macrophage-mediated gemcitabine resistance, we set out to develop an assay to functionally assess the response of ODCs to gemcitabine in the presence and absence of macrophages, and with or without a hypoxic gradient.

To investigate the chemoprotective effect of macrophages on ODCs in normoxia, we compared the dose response curves of gemcitabine-treated single-layer TRACER ODC monocultures and co-cultures using the alamarBlue assay. alamarBlue is a non-fluorescent resazurin dye that is reduced by metabolically active cells into a fluorescent compound that can be detected by typical fluorescent plate readers. Instead of a single cell-based readout such as flow cytometry, we chose alamarBlue because it is quick, simple, and can be used without digesting cells out of TRACER, which could compromise cell viability readouts that rely on intact cell membranes. Since alamarBlue provides a bulk measurement of cell viability, we first confirmed that the dominant alamarBlue signal obtained was from the ODCs and not the less metabolically active macrophages (**Figure S6a).** Having addressed this technical concern, we then treated single-layer, non-rolled ODC monocultures and co-cultures with varying concentrations of gemcitabine for three days (72 hours) to generate dose response curves for each condition (**Figure 4a**). When we measured the viability of each treated sample relative to their corresponding untreated controls in single layers, we observed a shift in the dose response curves for ODCs in co-culture upwards from the monoculture curves (**Figure 4b**). Correspondingly, ODCs in co-culture had a higher IC50 compared to the ODC monocultures (192 µM for the co-culture versus 98 µM for the monoculture) (**Figure 4b**), suggesting that the human-derived macrophages in our system exerted a chemoprotective effect on patient ODCs, similar to the observations in studies using mouse models^93,94^. Next, we assessed the effect of macrophages on gemcitabine resistance in a hypoxic gradient (rolled TRACERs) by treating 24-hour rolled ODC monocultures and macrophage-ODC TRACER co-cultures with either 98µM gemcitabine, the previously determined IC50 for ODCs in normoxia (single-layer TRACERs), or DMSO (vehicle control) for three days (72 hours) before unrolling for layer-specific analysis using the alamarBlue assay; cell viability was reported relative to each corresponding layer in the DMSO control-treated TRACERs (**Figure 4c**). To confirm that the DMSO concentration used was not lethal to either cell type, we also treated ODCs and macrophages with the DMSO concentration equivalent to 98 µM of gemcitabine (∼0.98% DMSO) (**Figure S6b)**. Interestingly, we found that the rolled macrophage-ODC TRACER co-cultures showed a significantly graded resistance to gemcitabine (slope of regression line different from 0: p-value = 0.0244), a phenomenon not observed in the rolled ODC TRACER monocultures in the culture media used for these studies (**Figure 4bii**). Together, these data suggested that in our system, gemcitabine resistance of ODCs is mediated by the presence of macrophages and that this macrophage-mediated resistance or chemoprotective effect could be exacerbated by hypoxic gradients.

Contrary to our group’s previous observation that rolled ODC monocultures grown in organoid CGM experienced a graded resistance to gemcitabine^35^, we reported in this study that rolled ODC monocultures in RPMI-1640 (complete) did not experience any graded response (**Figure 4bii)**. Again, we expect that this can be attributed to media differences, wherein the more basal-like phenotype induced by the ODCs in RPMI-1640 (complete) could have reprogrammed the cells to a more chemo-resistant phenotype at baseline normoxia which diminished any impact hypoxia might have on drug resistance. Since basal PDAC is associated with more aggressive disease, this further implied that the TRACER co-culture system described here could be important for understanding response to chemotherapy in hypoxic, basal disease. However, while there has been previous evidence to suggest that hypoxia drives resistance to chemotherapies^95–97^ and is associated with basal PDAC at the transcriptional level^98^, to our knowledge, this is the first study to show the impact of both macrophages and hypoxia on gemcitabine resistance in human pancreatic cancer cells. Further investigation of these important cell-cell interactions perhaps by combining TRACER with more granular analysis techniques such as RNA-sequencing is warranted and could have significant implications in understanding the mechanisms of how hypoxia functionally reprograms macrophage-cancer interactions, especially in basal-like PDAC, to contribute to therapy resistance, a metric that holds clinical relevance in the drug discovery pipeline.

**Figure 3:**
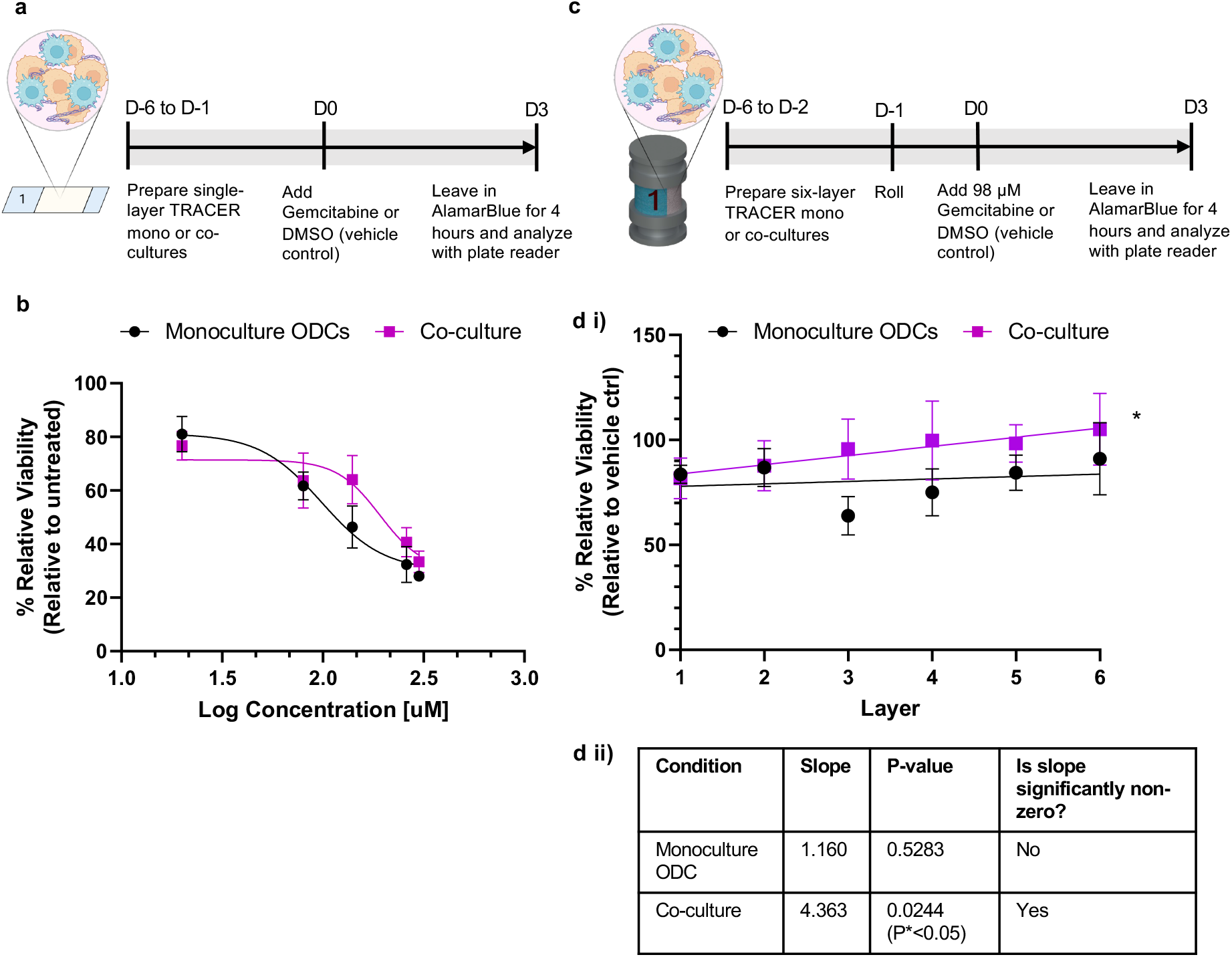
Response to the chemotherapy Gemcitabine in single-layer and rolled ODCs in co-culture with macrophages. **(Ai)** Schematic of the workflow for evaluating response to Gemcitabine in single-layer ODC co-cultures versus monocultures. The metabolic assay alamarBlue was used to analyze cell viability. **(Aii)** Dose response curves of Gemcitabine-treated single-layer ODC monocultures of ODC-macrophage co-cultures. Cell viability is expressed relative to untreated controls in either ODC monoculture or co-culture. A 4-parameter logistic curve was fit to the data to obtain IC50 values (ODC monoculture IC50 = 98 µM; ODC co-culture IC50 =198 µM). **(Bi)** Schematic of the workflow for evaluating response to Gemcitabine in rolled ODC co-cultures versus monocultures. After 24h of rolling, TRACER constructs were treated with either Gemcitabine (98 µM) or vehicle DMSO (0.98%) for 72 hours (3 days). **(Bii)** Viability of rolled ODC co-cultures and ODC monoculture TRACERs, 72 hours after treatment. Viability was calculated relative to each corresponding layer in the vehicle (DMSO)-treated TRACERs (i.e., Layer 1 gemcitabine treated co-culture TRACER is normalized to Layer 1 DMSO-treated co-culture) (Slope of best fit line vs. 0: p*<0.05). Error bars are mean ± SEM for n= 3-4 biological replicates.

### 3.7 Secreted factors produced by macrophages and ODCs in the inner layers of TRACER attenuate the T-cell inflammatory response

Having established that macrophages appeared to be functionally reprogrammed, at least to some extent, by hypoxia to exert a chemoprotective effect on ODCs, we next set out to determine if macrophage-ODC interactions were also rewired to impact their influence on T-cell function. In various cancers including PDAC, hypoxia and TAMs have separately been shown to impede T-cell effector function by indirectly secreting molecules such as cytokines and proteins that suppress T-cell response^99^. Thus, as proof-of-concept, we cultured primary T-cells with conditioned media retrieved from various TRACER layers to probe the indirect impact of hypoxia reprogrammed macrophage-cancer interactions on T-cells. First, we generated conditioned media by incubating layers one (normoxia), three (moderate hypoxia) and six (severe hypoxia) for 18 hours (after 24-hour of rolled culture) from TRACER macrophage monocultures, ODC monocultures, and macrophage-ODC co-cultures (**Figure 5a**). As a positive control, we also generated 18-hour conditioned media from cells cultured in a 0.2% hypoxia chamber for 24 hours. We then treated PBMC-derived T-cells with these conditioned medias collected from each layer (50% conditioned media, 50% fresh RPMI-1640 complete) for 24 hours to expose T-cells to any secreted factors from the TRACER constructs. Next, we activated T-cells with anti-CD3/CD28 for another 48 hours and finally, performed flow cytometry analysis (**Figure 5a**) to assess the inflammatory response of the exposed T-cells. Activating T-cells with anti-CD3/CD28 was essential for T-cells to proliferate and expand *in vitro*, elicit an inflammatory response, and mimic the *in vivo* situation wherein T-cells are usually already activated when in the tumour microenvironment^100–103^. To assess T-cell inflammatory response, we measured the percentage of T-cells that expressed the inflammatory molecules tumour necrosis factor alpha (TNFα) and Granzyme B (GZMB), which are critical effector function molecules in both CD8^+^ and CD4^+^ T-cells^100,104–107^. When we assessed CD8^+^ T-cells, the population of T-cells mainly responsible for exerting a cytotoxic function^108^, we observed a significant decrease in the % TNFα^+^ (L1 vs L6: p value = 0.0145; L1 vs 0.2% hypoxia: p-value = 0.0185) and % GZMB^+^ (L1 vs. L6: p-value = 0.0454;L1 vs 0.2% hypoxia: p-value = 0.0003) CD8 T-cells in layer six and in the 0.2% Hypoxia controls but only in co-cultures (**Figure 5b**). Correspondingly, we observed a decrease in double positive TNFα^+^GZMB^+^ CD8^+^ T-cells only in layer six and the positive hypoxia controls only for co-cultures (L1 vs. L6: p-value = 0.009; L1 vs 0.2% hypoxia: p-value = 0.0007) (**Figure 5b**); no change was observed in the CD8 T-cells treated with conditioned media from either macrophage or ODC monocultures. These results suggest that both macrophage and tumour cells, and potentially their interactions, are required to suppress the expression of pro-inflammatory proteins like TNFα and GZMB in CD8^+^ T-cells, and that this attenuation is likely hypoxia dependent.

We also assessed the impact of the conditioned media on CD4^+^ T-cells, the population of T-cells mainly responsible for exerting helper functions^109^, and observed a significant decrease in %TNFα^+^CD4^+^ T-cells treated with conditioned media collected from layers three (L1 vs L3: p value = 0.0218) and six (L1 vs L6: p value = 0.0009) of the TRACER co-cultures. Correspondingly, we also observed a decrease in the proportion of double positive %TNFα^+^ GZMB^+^ in CD4^+^ T-cells (L1 vs L6: p value = 0.0433) treated with conditioned media from layer six of the TRACER co-cultures (**Figure 5c**). Similar to our observations in CD8 T-cells, we did not observe a decrease in the proportion of %TNFα^+^, %GZMB^+^, or %TNFα^+^ GZMB^+^ CD4^+^ T-cells when they were exposed to conditioned media generated from either macrophage or ODC monoculture TRACERs. This further supports the idea that both macrophage and cancer cells, and potentially their interactions, in the inner layers of TRACER produce secreted molecules that sufficiently attenuate the inflammatory response in CD4^+^ T-cells. Further, given the observed decrease in %GZMB^+^ (L1 vs 0.2% Hypoxia: p value = 0.0262), %TNFα^+^ (L1 vs 0.2% Hypoxia: p value = 0.0018) and %GZMB^+^ TNFα^+^ (L1 vs 0.2% Hypoxia: p value = 0.0186) CD4^+^ T-cells in the 0.2% hypoxia co-culture controls, the diminished CD4 T-cell inflammatory response was likely dependent on hypoxia-driven macrophage-tumour phenotypes (**Figure 5c**). Intriguingly, we also observed a significantly increased proportion of GZMB^+^ CD4 T-cells in T-cells treated with conditioned media from layer six (L1 vs L6: p value = 0.0179) and from the positive hypoxia control (L1 vs 0.2% Hypoxia: p value = 0.0273) of monoculture macrophage TRACERs (**Figure 5c**). While macrophage TRACER monocultures did not enable the establishment of a measurable hypoxic gradient, at least during the timeframe of our experiments, it is possible that the macrophages in these cultures did generate inflammatory signals in response to other stresses which in turn resulted in conditioned media with the capacity to elevate GZMB expression in the CD4+ T-cells. The resulting increased expression of GZMB in CD4 T-cells could potentially impact the TME in several ways, including increasing cytolytic activity against CD8 T-cells^110^ or enhancing cancer cell response to checkpoint inhibitors^111^. A deeper analysis of CD4 T-cell subsets and the conditioned media secretome would potentially be useful therefore in the future to explore the mechanism of this GZMB increase.

One concern during our analysis was that decreases in CD8 and CD4 T-cell inflammatory molecule expression levels were due to macrophage density differences between the different samples, that ultimately could produce changes in the secreted cytokine levels present in the conditioned media. Using confocal microscopy, we confirmed that layer six TRACERs had similar numbers of cells compared to the other TRACER layers (**Figure S7a**). Further, to confirm that any minor differences in cell numbers between the different TRACER layers after the rolled culture periods did not result in different dilutions of secreted factors in the condition media, we assessed the proportions of inflammatory molecules TNFα and GZMB in T-cells treated with varying dilutions of conditioned media from layer six TRACER co-cultures and observed similar decreases (relative to layer 1) for all conditioned media solutions tested (**Figure S7bi-ii**). Since the proportion of TNFα^+^ and GZMB^+^ cells overall did not differ from each other, we concluded that the observed diminished inflammatory response in CD8 and CD4 T-cells were likely not due to any minor cell density differences between each layer. Of note, we also investigated the impact of conditioned media on another pleiotropic mediator of tumour inflammation, IFNγ^112^ but did not observe any significant change in expression for any of the conditions (**Figure S7ci-ii**). Additionally, we found that the percentage of IFNγ positive cells were noticeably low (<2%) in general, including the samples only stimulated with anti-CD3/CD28 (not treated with conditioned media) (**Figure S7d**). Since IFNγ is usually a temporally mediated early response to inflammation^112–114^, it is possible that we were not able capture the timepoint of IFNγ expression of both CD8 and CD4 T-cells. In future studies performing in-depth evaluations of T-cell response in TRACER co-cultures, it will likely be important to consider the temporal dependence of certain inflammatory response molecules.

Overall, our data support the idea that hypoxia-enriched tumour-macrophage interactions play a role in attenuating T-cell inflammatory response in our system. Further, our data highlight the utility of the TRACER co-culture system for functionally exploring how phenotypic changes in hypoxia-reprogrammed cells can modulate T-cell function, which could be useful for identifying and exploring the molecular mechanisms of novel immunotherapeutic targets.

**Figure 3.4:**
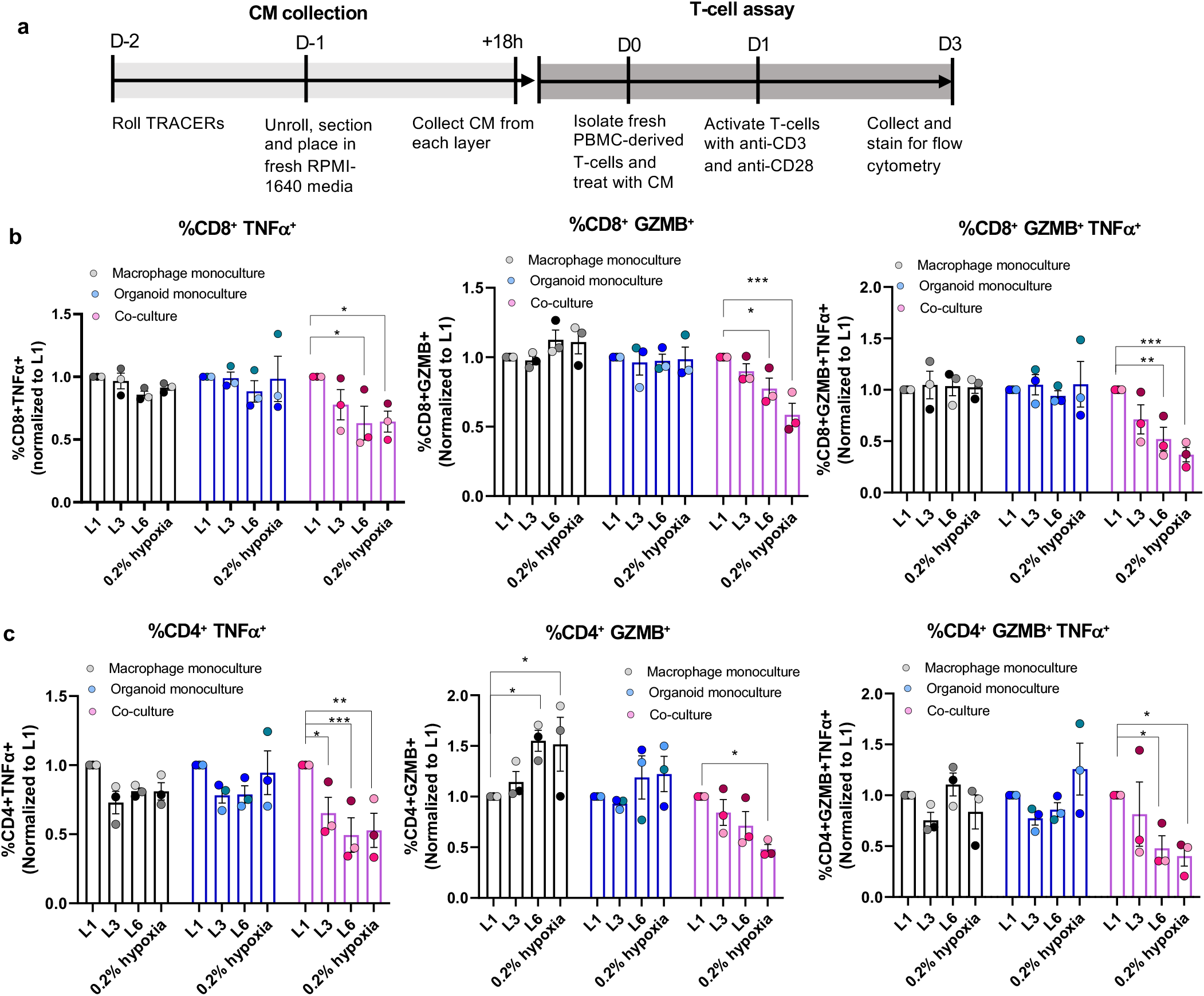
CD8+ and CD4+ PBMC-derived T-cells exposed to conditioned media from macrophage-ODC TRACER layers. **(A)** Schematic of experimental design for collecting conditioned media and treating T-cells with conditioned media. 18h conditioned media were collected from Layers 1, 3 and 6 of 24h rolled macrophage monoculture, ODC monoculture and macrophage-ODC co-culture TRACERs and stored until ready for use. Freshly isolated PBMC-derived T-cells were treated with layer-specific conditioned media for 24h, activated with anti-CD3/CD28 for another 48 hours and then collected for analysis by flow cytometry. **(B)** Quantification of CD8 T-cells treated with conditioned media from various TRACER layers, including positive hypoxia controls. %CD8^+^TNFα^+^, %CD8^+^GZMB^+^ and %CD8^+^TNFα^+^GZMB^+^ T-cells were normalized to Layer one in each condition (p*<0.05, p**<0.01, p***<0.01). **(C)** Quantification of CD4 T-cells treated with conditioned media from various TRACER layers, including positive hypoxia controls. %CD4^+^TNFα^+^, %CD4^+^GZMB^+^ and %CD4^+^TNFα^+^GZMB^+^ T-cells were normalized to Layer one in each condition (p*<0.05, p**<0.01, p***<0.01). Representative flow cytometry plots and a representative gating strategy is shown in **Figure S8a-b**. Error bars are mean ± SEM for n= 3 biological replicates. Each dot on the bar graph represents one independent T-cell donor treated with conditioned media from 3 independent TRACERs.

## 4. Discussion

The role of the highly dysregulated PDAC TME in contributing to disease progression is well-known. Among the components of the PDAC TME, hypoxia is a prominent gradient that has been shown to reprogram cancer cells to confer a survival advantage, while macrophages are present in high numbers in the PDAC TME and have been linked to poor prognosis and chemoprotective and immune-regulatory phenotypes^5,9–11,115,116^. However, deciphering complex interactions between these prominent components of the PDAC TME is challenging in part due to the lack of available culture models that allow the contribution of each factor and their interactions to be systematically studied. Specifically, there are currently no fully human *in vitro models* that simultaneously incorporate macrophages, cancer cells and hypoxia, are physiologically relevant and incorporate patient cells, and allow for the exploration of these intersecting parameters. Here, to address this gap, we have developed a fully human 3D *in vitro* model containing primary macrophages and patient organoid cells that enables systematic interrogation of cancer-immune cell interactions in a cell generated hypoxic gradient.

Our group previously developed an engineered paper-based 3D in vitro model called TRACER that enables culture of patient ODCs to establish cell-derived hypoxic gradients^21,33,35^. In this study, we first assessed the feasibility of culturing primary macrophages in TRACER and confirmed that they retained expected plasticity to pro and anti-inflammatory phenotypes when cultured in the TRACER scaffold. Since macrophages are known to be highly plastic cells, we were not surprised that macrophage activation due to external stimuli was still detectable over any baseline cell activation resulting from the biomaterial scaffold. However, because macrophages experienced some activation from the TRACER scaffold, culturing macrophages as monocultures in TRACER without any exposure to external stimuli (e.g., activating cytokines, co-culture with another cell type) should be interpreted cautiously. Nevertheless, here we show that culturing primary macrophages in scaffold-supported cultures such as TRACER is possible and importantly, could be integrated into other similar models with different applications, such as the SPOT model developed by our group for high-throughput cell characterization^117,118^ or those developed by the Whitesides and Lockett group^119,120^.

We invested significant efforts to optimize media conditions for co-culturing ODCs and macrophages together and identified a ODC pre-conditioning strategy to maintain viability, maturation, and polarity in ODCs while also ensuring co-cultures could be established in a media free of organoid media growth factors, which were shown to prematurely activate macrophages. While our specific media strategy was developed for macrophages and ODCs, we speculate the blending approach we have developed could be extended to the co-culture of ODCs with other immune cells such as T-cells, myeloid derived suppressor cells (MDSCs), natural killer (NK) cells, dendritic cells (DCs) or engineered immune cells which are all typically cultured in RPMI-1640^121–125^. Of note, ODCs cultured using our media blend strategy exhibited a phenotype resembling basal-like PDAC. This was consistent with results from previous studies which reported on the plasticity of organoids in various media^53^. Although, we did not investigate this switch to a more basal phenotype in depth here, our results suggest that our TRACER co-culture platform could be used to specifically model basal disease, the PDAC subtype associated with poorer prognosis and chemoresistance, for which no *in vitro* models for exploring TAM behaviour currently exist. We also noted dynamic changes in the behaviour of the different cell populations over time and determined that 24 hours of co-culture provided sufficient time to observe expected phenotypic changes in co-cultures. Selecting one specific time point was necessary because TRACER analysis is typically destructive using endpoint assays. Given that macrophage-cancer interactions are dynamic, the need to select one or two timepoints for analysis is a limitation of the TRACER platform and future work to exploring macrophage-tumour interaction dynamics will likely require the use of different models compatible with continuous or longitudinal studies, such as the GLAnCE platform developed in our group^126,127^. For example, live-cell assays that track macrophage phagocytotic capacity or antigen presentation could add to our knowledge of macrophage function in the TME. Despite these limitations, TRACER remains uniquely positioned to probe the impact of tumour-macrophage interaction in cell-generated microenvironmental gradients in a spatially resolved manner and potentially for basal-like PDAC, which has thus far not been demonstrated with other engineered models.

We also invested significant effort to validate our co-culture model by confirming the presence of hypoxic gradients in TRACER cultures and demonstrating a number of known biological responses to hypoxia that have been previously shown to occur in PDAC tumour cells and macrophages. However, we also note that the expected transcriptional response for some genes we measured were not observed, specifically in macrophages co-cultured with ODCs in TRACER (IL-6 and IL-1β) (**Figure S4c**). This was likely due to a combination of factors such as donor variation from using primary cells, heterogeneity of TAM phenotype, and the challenges of reconciling expected patterns from published studies, that mainly focus on mouse-macrophages, with observations in our study, that used human-macrophages. Moreover, while several studies focus on hypoxia, very few explore the impact of microenvironmental gradients (e.g., nutrients, metabolites, and waste) on human macrophages. Thus, in addition to heterogeneities in TAM phenotype, interpreting patterns of macrophage response in TRACER, wherein both hypoxic and molecular gradients are incorporated, is even more challenging. Here, we were restricted to investigating only a few genes using qPCR; therefore, to resolve some of these challenges, future studies that apply single cell or omics technologies with TRACER could lead to the exploration of phenotypic changes in macrophage-ODC interactions with more granularity. Some examples of these future studies could involve investigating macrophage heterogeneity in hypoxia and other relevant microenvironmental gradients in the TME, identifying what pathways are significantly graded in both macrophages and cancer cells, and importantly, linking observations in TRACER with patient data to validate the utility of our model in recapitulating patient tissues.

Aside from validating known biology in the TRACER co-culture, we demonstrated the utility of TRACER for gaining novel insights about tumour-macrophage interactions in a graded microenvironment. One example of this is showing TRACER’s capacity to distinguish between the impact of hypoxia versus other cell-derived gradients by comparing the expression of certain genes in TRACER cultures versus single layer TRACER cultures placed in a hypoxia chamber. Although we did not define these other gradients or deeply explore their influence on cancer-macrophage interactions here, TRACER has been used previously to define and explore such gradients (e.g., small molecules and metabolites)^33^, and exploring how metabolic gradients impact cancer-immune interactions, for example, could be an exciting area for future study, given that there is high metabolic heterogeneity in the TME^128,129^. Another example to highlight the utility of TRACER is by demonstrating the potential clinical relevance of TRACER for evaluating response to therapy. Here, we found that macrophages exerted a chemoprotective effect on ODCs in single layer, normoxic co-cultures, which was consistent with previous studies in cell lines^94^ and mouse models^93^. Excitingly, we showed in a fully human model that ODCs in co-culture with macrophages in TRACER displayed a significantly graded resistance to gemcitabine but not in monoculture TRACERs, suggesting that hypoxia and other gradients generated in TRACER could have rewired macrophage-ODC interactions to promote chemotherapy resistance. Although our study was limited to measuring bulk responses to gemcitabine treatment in TRACER, with more granular analysis, we could potentially also investigate the mechanisms of chemoprotection and resistance to gemcitabine (or perhaps other clinically relevant drugs) between macrophages and ODCs in more detail.

Finally, in addition to evaluating response to therapy, we showcased the capacity to generate conditioned media from TRACER layers, which presented a method to interrogate how secreted factors produced by cells in various layers affect T-cell response, without having to set-up more complicated systems such as tri-cultures. In this study, we found that CD8 and CD4 T-cells treated with conditioned media collected from the inner layers (layer six) of TRACER co-cultures, but not in ODC or macrophage monocultures, decreased the percentage of inflammatory TNFα^+^, GZMB^+^ and TNFα^+^GZMB^+^ T-cells in a hypoxia-dependent manner. Although the exact mechanisms of how macrophage-cancer cell interactions are rewired in TRACER were not investigated, the results presented here suggested that interactions between macrophages and cancer cells could potentially be modulated for therapeutic benefit. Future work could extend to evaluating how secreted factors induce functional changes in cytotoxic T-cell capacity, T-cell subtypes (e.g., exhausted, effector-memory, regulatory, etc.), or even in other TME components such as CAFs. Thus, identifying what these secreted molecules are (potentially using omics technologies) and understanding how they are produced could have important implications in deciphering the interactions between macrophages and pancreatic cancer cells in hypoxia.

## 5. Conclusion

Here we describe a TRACER co-culture system that provides a physiologically relevant, fully human 3D in vitro model of PDAC tumour-macrophage interactions in the TME. This tool is particularly useful for deciphering the responses of macrophages and cancer cells in hypoxia and other chemical and molecular gradients, which is challenging to evaluate using existing models. In this study, we showed that co-culturing macrophages with ODCs was feasible. Additionally, we optimized media conditions, developed co-culture TRACER protocols, and confirmed that macrophages and ODCs experienced graded levels of hypoxia and expressed expected responses to hypoxia while rolled in TRACER. Interestingly, profiles of graded gene expression (TGFβ and PD-L1) varied between ODC TRACER monocultures and co-cultures, indicating that hypoxic macrophages exerted a significant influence on ODCs to promote immune-regulatory cancer cell phenotypes. Moreover, we established a functional assay to evaluate response to gemcitabine and found that macrophages exerted a chemo-protective effect on ODCs in single-layer (normoxic) TRACER layers and in rolled (hypoxic) TRACER co-cultures, showing for the first time in a human model of PDAC that hypoxia-reprogrammed macrophages exacerbated resistance to gemcitabine. Finally, we demonstrated that hypoxia-reprogramming of macrophage-tumour interactions in the inner layers of TRACER indirectly suppressed T-cell inflammatory response, suggesting that targeting hypoxic macrophage-cancer interactions could be an effective therapeutic strategy to improve PDAC response to immunotherapies. We anticipate that this work will enable a unique mechanistic investigation of primary, human, cancer-macrophage interactions as influenced by the hypoxic TME, which overall, can help explain why current therapies fail in PDAC and inform the development of novel therapies.

## Acknowledgements

This work was funded by an Ontario Graduate Scholarship to ILC and JLC, a Vanier Canada Graduate Scholarship to NLB, a Canada Institute of Health (CIHR) Research Project Grant to APM.

