## Supplementary Information for "An Engineered 3D Co-culture Model of Primary Macrophages and Patient-Derived Tumour Cells to Explore Cellular Responses in the Graded Hypoxic Microenvironment of Pancreatic Cancer"

### Supplementary Figures

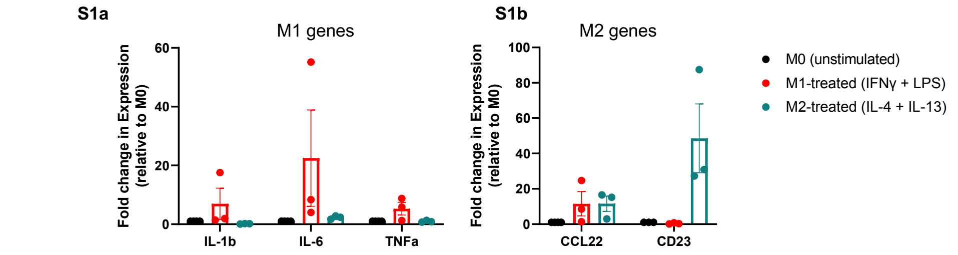


**Figure S1:Gene expression of macrophages in gel only configuration**. **(A)** qPCR analysis of pro-inflammatory (M1) genes IL-1b, IL-6 and TNFα in macrophages treated with either M0 (unstimulated), M1 (IFNγ + LPS), or M2 (IL-4 + IL-13). **(B)** qPCR analysis of pro-inflammatory (M2) genes CCL22 and CD23 in macrophages treated with either M0 (unstimulated), M1 (IFNγ + LPS), or M2 (IL-4 + IL-13). For all conditions, gene expression was quantified relative to the housekeeping gene 18S and normalized to the expression of M0 (unstimulated) macrophages and expressed as a fold change (ΔΔCt). No statistically significant expression was observed. Error bars are mean ± SEM for n= 3 independent donors (biological replicates).

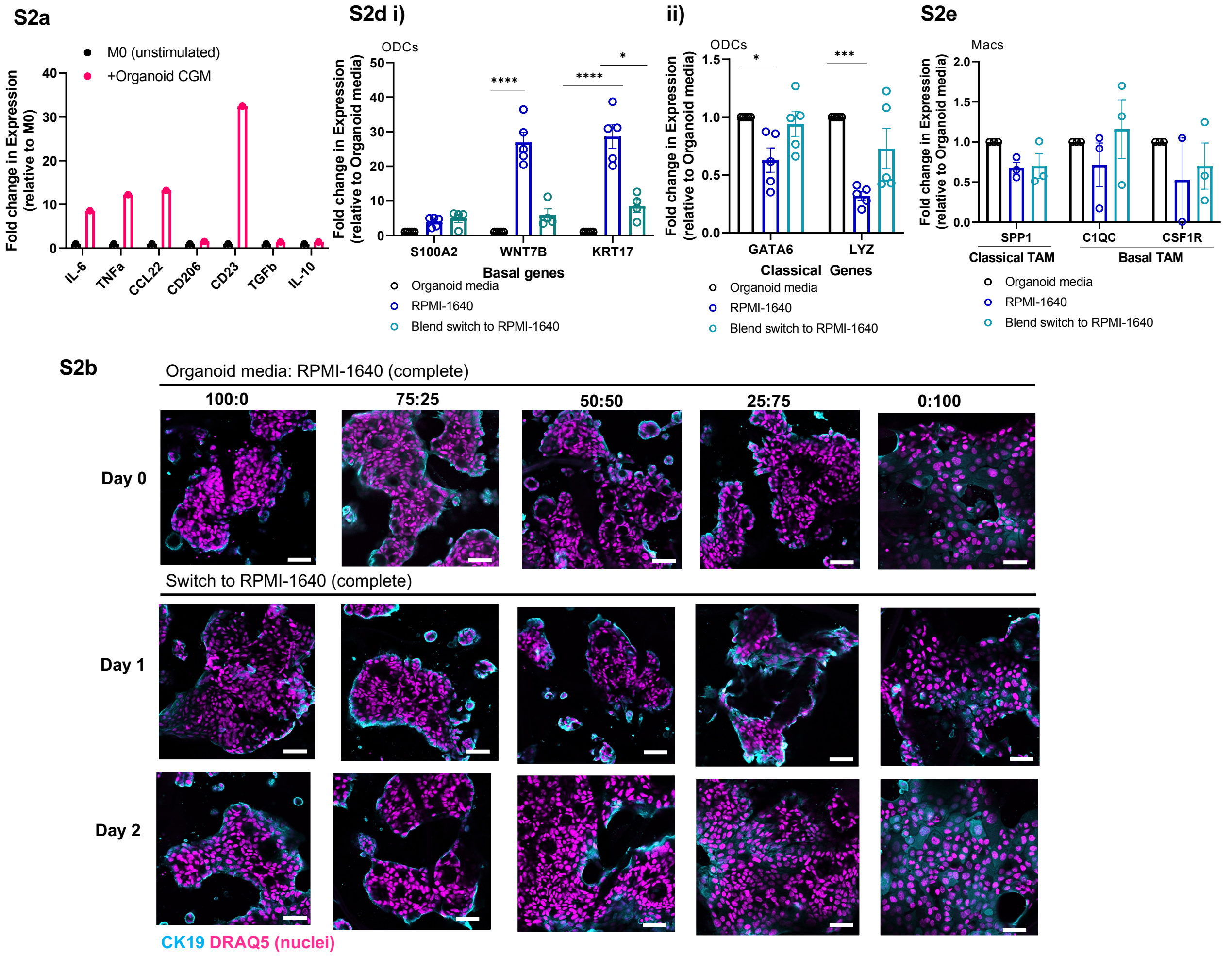

S2c

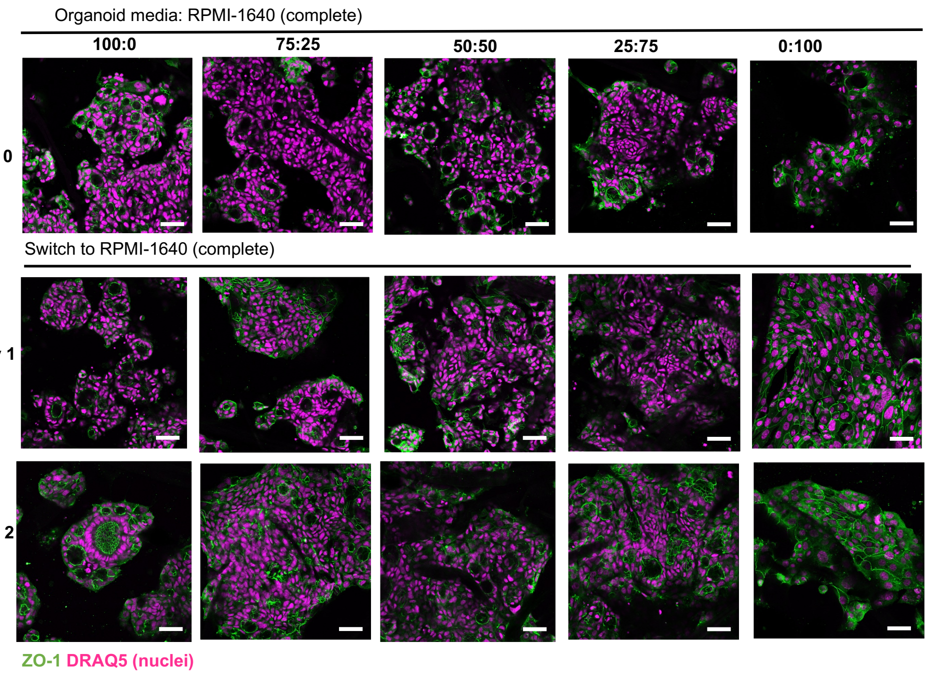


**Figure S2: The effect of media on macrophages and organoids. (A)** qPCR analysis of pro and anti-inflammatory genes expressed by macrophages activated with PDAC organoid growth media for 24 hours. Gene expression was normalized to the housekeeping gene GAPDH and expressed as a fold change relative to M0 (unstimulated) macrophages. N=1 biological replicate, 2 technical replicates. **(B)** Representative confocal images of CK19 expression of ODCs grown in various media regimes in single-layer TRACER scaffolds at Day 0, 1 and 2. Scale bar = 50 µm. **(C)** Representative confocal images of ZO-1 expression of ODCs grown in various media regimes in single-layer TRACER scaffolds at Day 0, 1 and 2. Scale bar = 50 µm. **(Di)** qPCR analysis of genes associated with the basal PDAC subtype (S100A2, WNT7B, KRT17) of ODCs in various media regimes in single-layer TRACER scaffolds for 6 days. Gene expression was normalized to the housekeeping gene RPLP0 and expressed as a fold-change relative to ODCs in PDAC organoid growth media (p*<0.05, p****<0.0001). **(Dii)** qPCR analysis of genes associated with the classical PDAC subtype (GATA6, LYZ) of ODCs in various media regimes in single-layer TRACER scaffolds for 6 days. Gene expression was normalized to the housekeeping gene RPLP0 and expressed as a fold-change relative to ODCs in PDAC organoid growth media (p*<0.05, p****<0.0001). **(E)** qPCR analysis of genes associated with the classical (SPP1) and basal (C1QC, CSF1R) PDAC TAMs of macrophages treated for 48-hours with conditioned media generated by ODCs grown in either organoid growth media, RPMI-1640 (complete) or the media blend strategy (75:25 organoid CGM:RPMI-1640) in domes. Gene expression was normalized to the housekeeping gene 18S and expressed as a fold-change relative to macrophages in PDAC organoid growth media (p*<0.05, p***<0.001). Error bars are mean ± SEM for n= 3 biological replicates,2-3 technical replicates (for ODCs) or n= 3 independent donors (for macrophages).

S3a

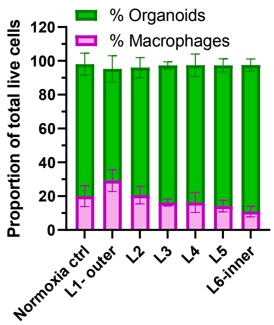


S3b

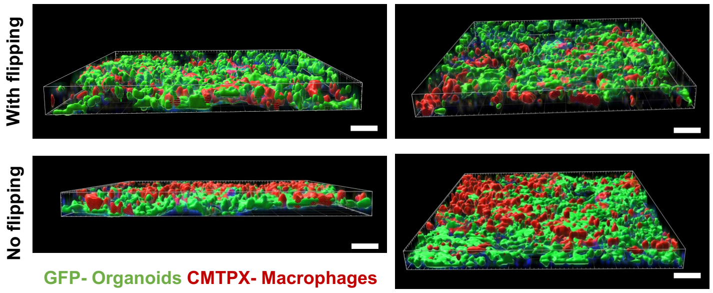


S3c

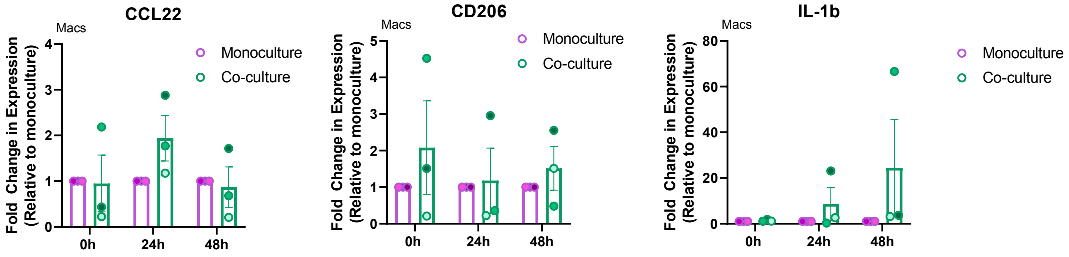

**Figure S3: Optimization of co-culture conditions for macrophages and ODCs. (A)** Proportions of live macrophages and organoids in single-layer TRACER scaffolds (Normoxia ctrl), and in each layer of the rolled TRACER scaffold. Proportions were measured using flow cytometry. Macrophages and ODCs were discriminated by CD45+ and CD45- signal. **(B)** Representative confocal images of GFP-ODCs co-cultured for 48h with CMTPX-labeled macrophages added onto the ODC-containing scaffolds with or without flipping. Scale bar = 80 µm**. (C)** qPCR analysis of CCL22, CD206 and IL-1b of macrophages in co-culture with ODCs versus monoculture macrophages at 0, 24h and 48h. The 0h timepoint represents the 24h after macrophages were added onto the ODC-containing scaffolds for co-culture. Gene expression was normalized to the housekeeping gene 18S and expressed as a fold change relative to monoculture macrophage controls. Error bars are mean ± SEM for n= 3 independent donors (for macrophages).

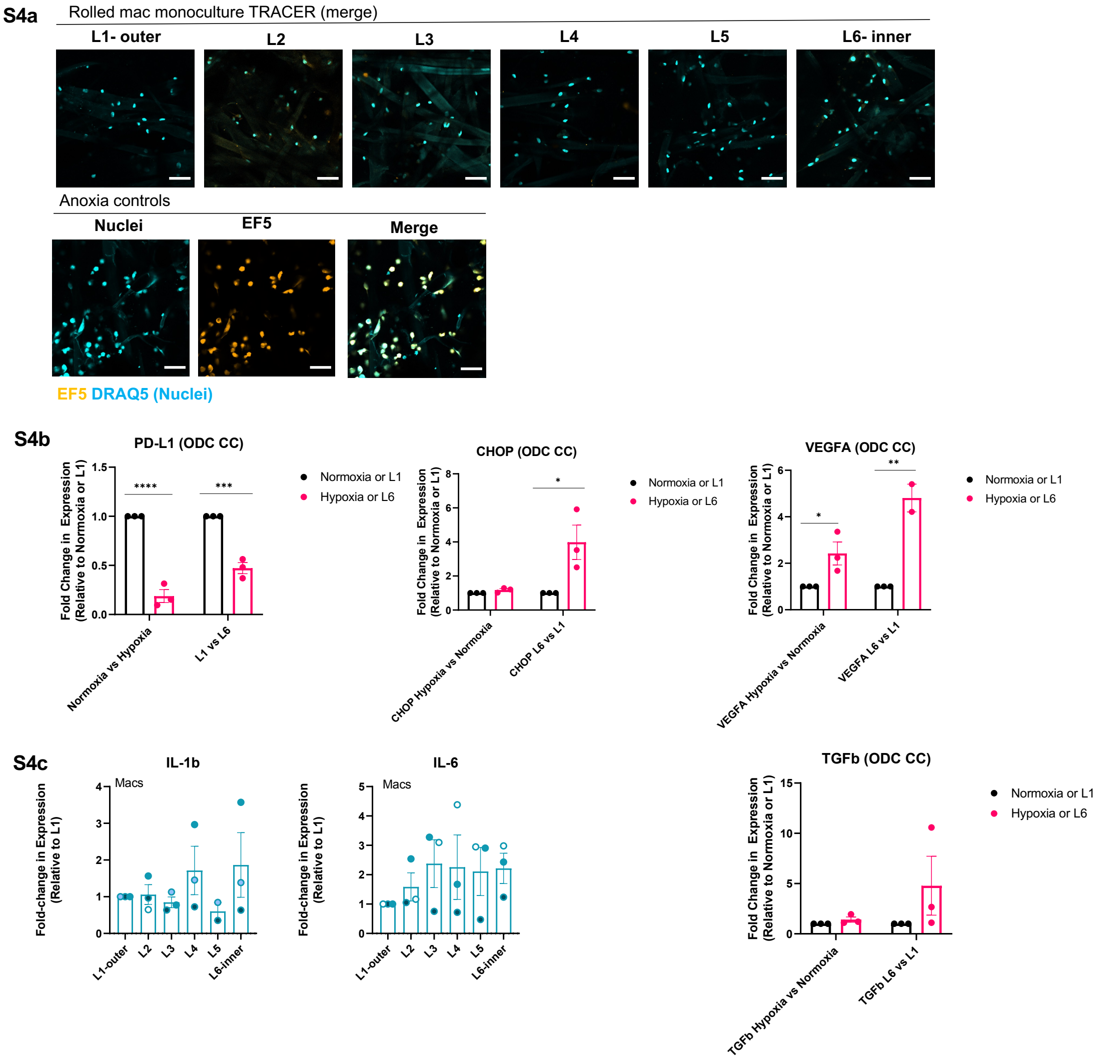


**Figure S4:Optimization of co-culture conditions for macrophages and ODCs in rolled TRACERs. (A)** Representative confocal images of EF5 expression in macrophage monoculture TRACERs and anoxia controls. Macrophage monocultures were either rolled for 24h in TRACER or left in an anoxic chamber (0% pO2) for 24h. Scale bar = 100 µm. **(B)** qPCR analysis of CHOP, VEGFA, TG and PD-L1 for ODCs in co-culture with macrophages in either TRACER or in single-layer scaffolds placed in 0.2% hypoxia chamber. Gene expression was normalized to the housekeeping gene RPLP0 and expressed as a fold change relative to either L1 (for TRACER) or normoxia (for 0.2% hypoxia controls). ($p^* < 0.05$, $p^{**} < 0.01$, $p^{***} < 0.001$, $p^{****} < 0.0001$). **(C)** qPCR analysis of IL-1b and IL-6 for macrophages in co-culture with ODCs in TRACER. Gene expression was normalized to the housekeeping gene 18S and expressed as a fold change relative to L1. Error bars are mean ± SEM for n= 3 biological replicates or n= 3 independent donors (for macrophages).

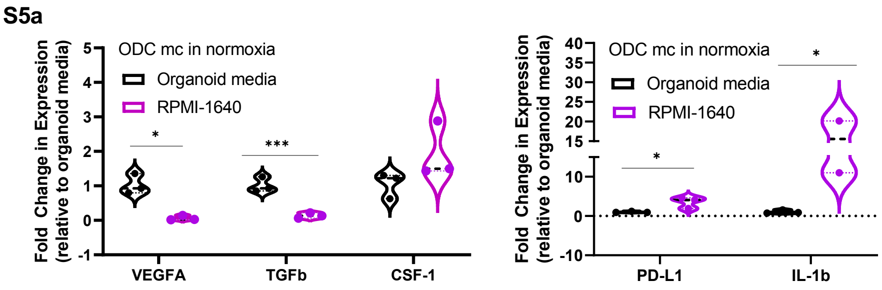


**Figure S5: Baseline gene expression of ODC monocultures in single-layer scaffolds (in normoxia). (A)** qPCR analysis of VEGFA, TGFb, CSF-1, PD-L1 and IL-1b in single-layer, normoxic ODC-monocultures cultured in either PDAC organoid complete growth media or RPMI-1640 (complete) for 6 days. Gene expression was normalized to the housekeeping gene RPLP0 and was expressed as a fold change relative to ODCs in PDAC organoid growth media (p*<0.05, p***<0.001). Error bars are mean ± SEM for n= 3 biological replicates.

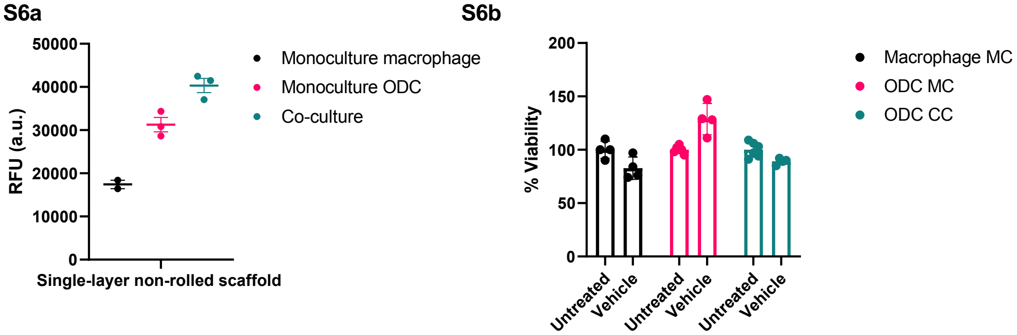


**Figure S6: Optimization of alamarBlue assay to evaluate response to gemcitabine in single-layer and rolled TRACERs. (A)** Relative fluorescence units (RFU) of untreated macrophage monoculture, ODC monoculture, and macrophage-ODC co-cultures in single-layer TRACER scaffolds incubated with alamarBlue (10v/v% alamarBlue to fresh RPMI-1640 media) for 4 hours. Error bars are mean ± SEM for n= 2-3 technical replicates. **(B)** Viability of macrophage monoculture, ODC monoculture and macrophage-ODC co-cultures in single-layer TRACER scaffolds treated with either nothing (untreated) or 0.98v/v% DMSO (vehicle control). 0.98% DMSO is the equivalent vehicle control for 98 µM gemcitabine, the dose used to treat rolled TRACERs and evaluate response to therapy. The viability is expressed as a percentage of the alamarBlue readings (RFU) obtained for vehicle control samples relative to the untreated samples. Error bars are mean ± SEM for n= 2 biological replicates, 2-3 technical replicates.

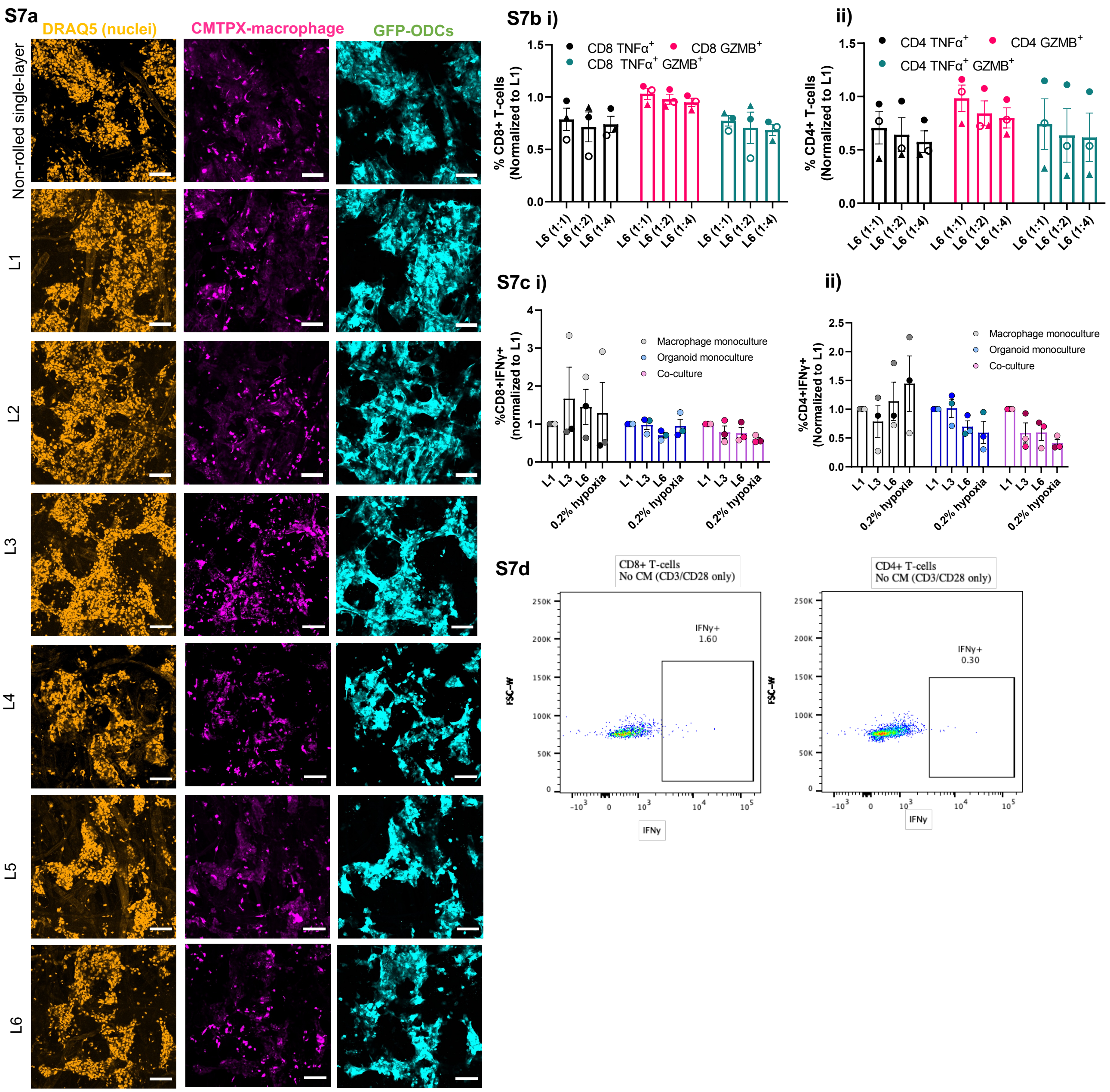


**Figure S7: Supplementary figure for T-cell conditioned media experiment. (A)** Representative confocal images of GFP-ODCs and CMTPX-labeled macrophages in 24h non-rolled single layer and rolled (layers one to six) TRACER co-cultures. Scale bar = 100µm. **(Bi)** Proportion of TNFα+, GZMB+ and TNFα+GZMB+ CD8 T-cells or **(Bii)** CD4 T-cells treated with CM collected from layer six TRACER co-cultures. Proportions were normalized to CD8 or CD4 T-cells treated with CM collected from layer one of TRACER co-cultures and acquired by flow cytometry. CM was used at varying dilutions with RPMI-1640 (complete) media (1:1, 1:2, 1:4). **(Ci)** Proportion of IFNγ+ CD8 T-cells and **(Cii)** CD4 T-cells treated with CM collected from macrophage monoculture, ODC monoculture or macrophage-ODC co-culture TRACERs and their respective hypoxia positive controls (CM collected from single-layer TRACERs placed in 0.2% hypoxia chambers). Proportions were normalized to CD8 or CD4 T-cells treated with CM collected from layer one of TRACER co-cultures and acquired by flow cytometry. **(D)** Representative flow cytometry plots of IFNγ+ CD8 and CD4 T-cells activated with anti-CD3/CD28 only (not treated with any CM). Error bars are mean ± SEM for n= 3 biological replicates (one independent T-cell donor, treated with CM from 3 independent TRACERs).

**S8a**

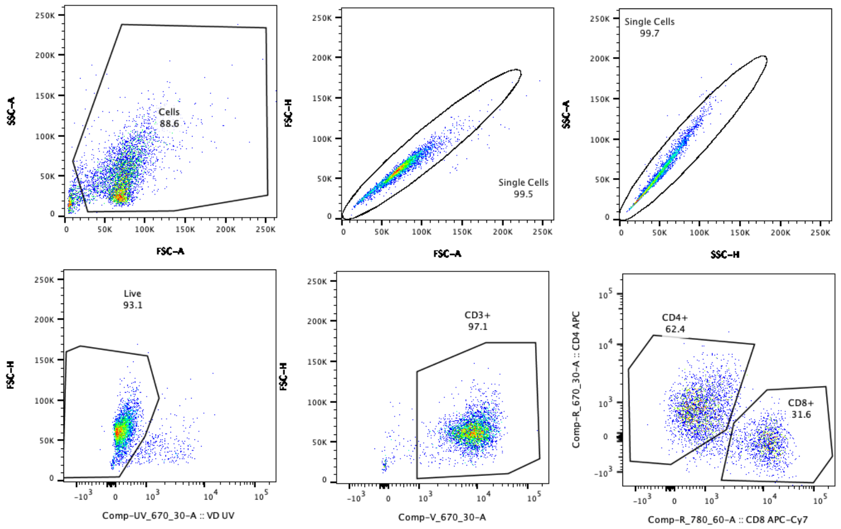


**S8b**

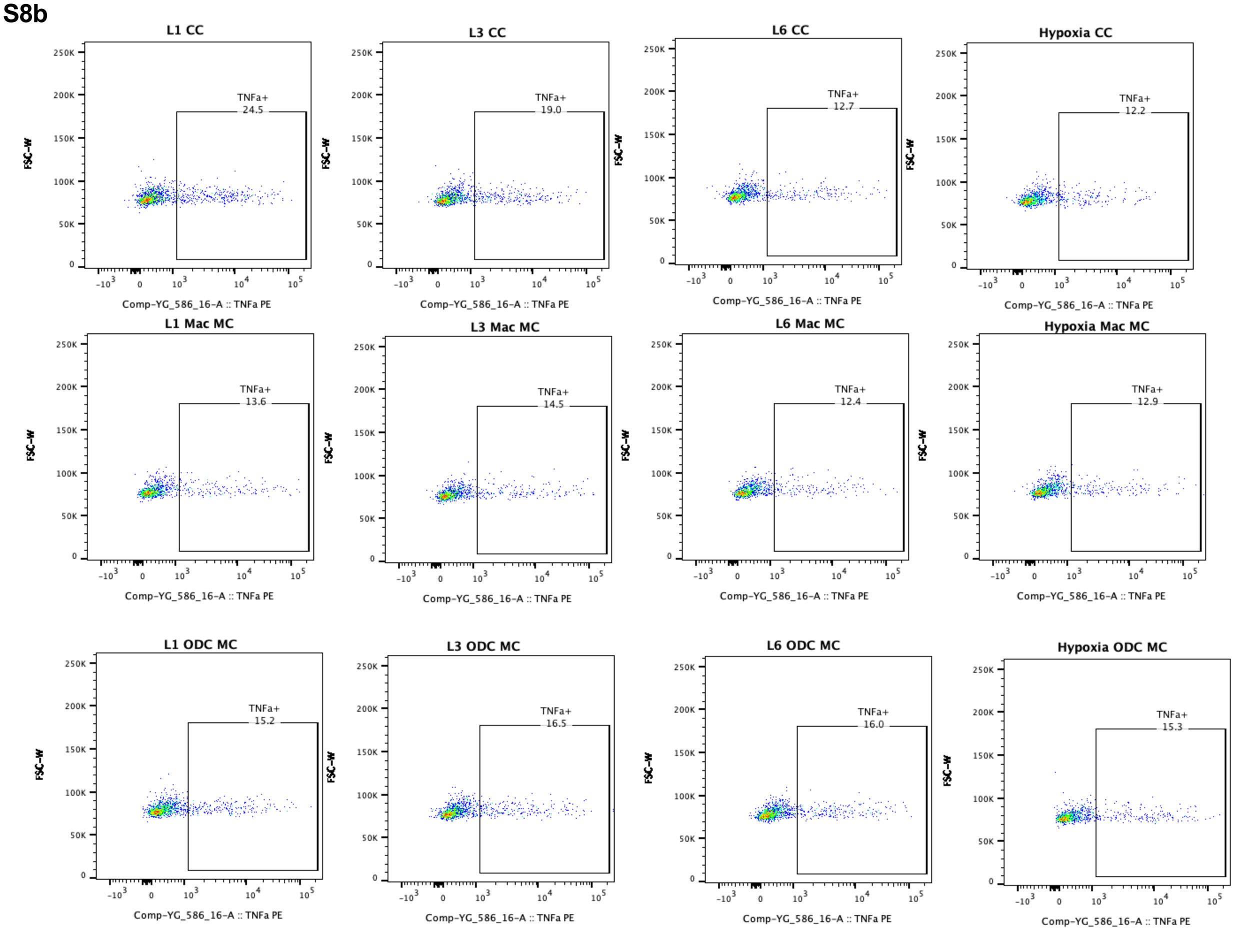


**Figure S8: Representative flow cytometry plots for the analysis of the T-cell conditioned media experiment. (A)** Representative flow cytometry plots showing gating strategy for analyzing T-cell flow cytometry data. Only live, single cells were analyzed; debris and doublets were excluded. CD3 was used as a pan T-cell marker and CD4 and CD8 T-cell populations were determined for further analysis in FlowJo. **(B)** Representative flow cytometry plots of TNFα+ CD8+ T-cells treated with CM collected from macrophage monoculture, ODC monoculture and ODC-macrophage co-cultures from layers one, three and six of rolled TRACER co-cultures. Flow cytometry plots of TNFα+ CD8+ T-cells treated with CM collected from single-layer TRACERs placed in a 0.2% hypoxia chamber were also included.

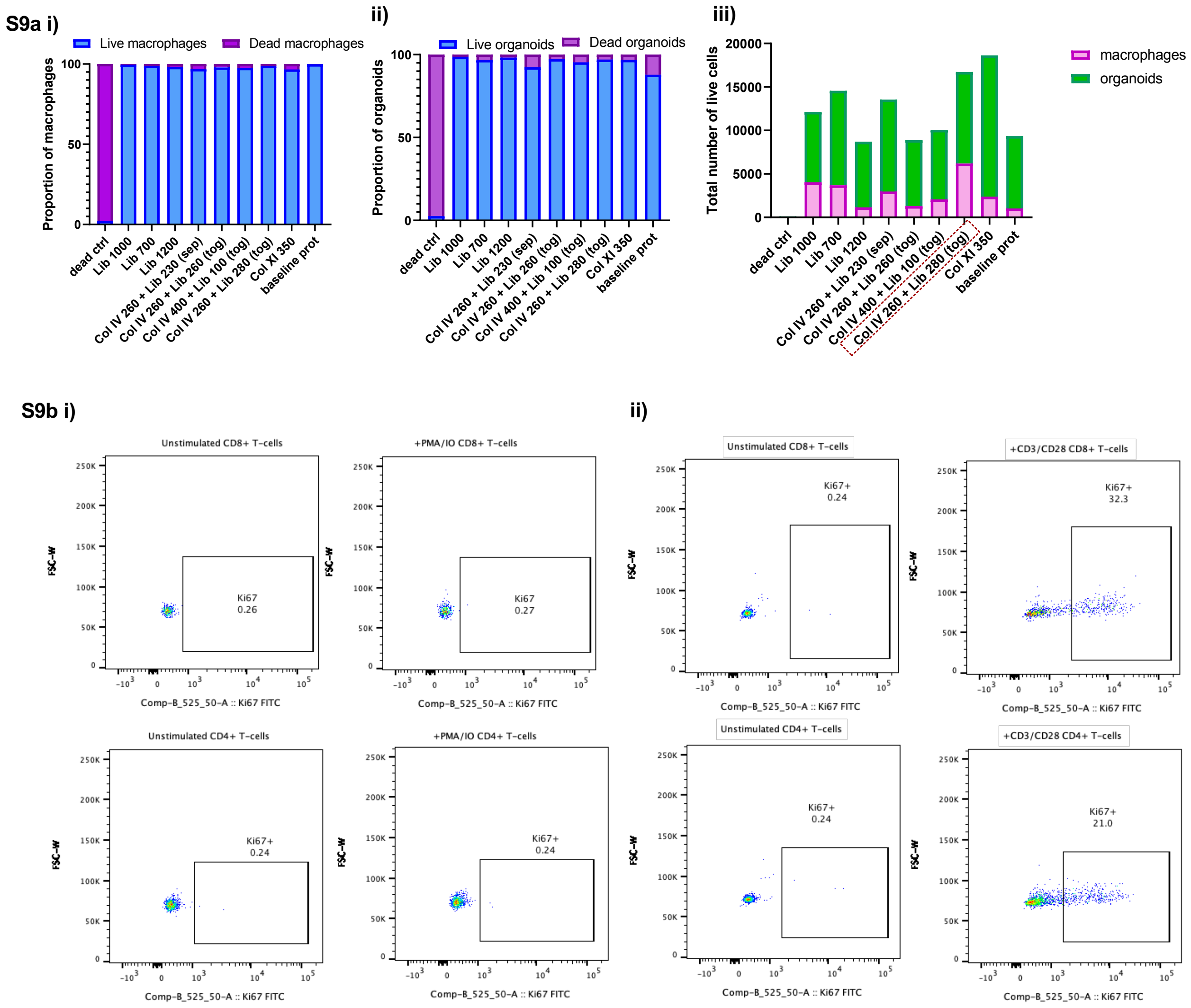


**Figure S9: Protocol optimizations. (A) Digestion protocol optimization.** Single-layer macrophage-ODC TRACERs were generated, and cells were subjected to various digestion protocols (Lib = Liberase TL; Col IV = Collagenase IV; Col XI = Collagenase XI; tog=together; sep= separate) and compared to a dead control (treated with 70% ice cold ethanol for 10 minutes) and the baseline digestion protocol used in the original TRACER study (cite Rodenhizer et al). All data was acquired using a flow cytometer. Live/dead cells were identified using Zombie Violet viability dye. Macrophages were discriminated from ODCs by CD45+ signal. **(Ai)** Proportion of macrophages digested out using various digestion protocols. Macrophage viabilities were similar across all conditions, except for the dead control. **(Aii)** Proportion of ODCs s digested out using various digestion protocols. ODC viabilities were similar across all conditions, except for the dead control. **(Aiii)** Total number of live macrophages and ODCs digested out of single-layer TRACER co-cultures. The greatest number of cells obtained was observed with the Col IV 260 + Lib 280 (tog) protocol (enclosed in a red box) and thus, was the protocol used moving forward. N=2 biological replicates, 2 technical replicates each. **(B) T-cell activation protocol optimization. (Bi)** Representative flow cytometry plots of Ki67+ (proliferation marker) CD8+ and CD4+ T-cells either unstimulated (media only) or stimulated with phorbol 12-myristate 13-acetate (PMA) and Ionomycin (IO). The proportion of Ki67+ CD8 and CD4 T-cells is similar between unstimulated and PMA/IO stimulated. **(Bii)** Representative flow cytometry plots of Ki67+ (proliferation marker) CD8+ and CD4+ T-cells either unstimulated (media only) or stimulated with anti-CD3/CD28. The proportion of Ki67+ CD8 and CD4 T-cells is around 20-30% for the stimulated T-cells and thus, we used the anti-CD3/CD28 protocol to activate T-cells.

**Table S1: Table of Statistics**

| Figure | n | Statistical test | P-values |
|---|---|---|---|
| **1Aii** | 3-6 independent donors, 3 technical replicates each | ANOVA with Tukey's multiple comparison post-hoc test | IL-1β: 2D vs Gel (p-value = 0.549); 2D vs. TRACER (p-value = 0.0008***); Gel vs. TRACER (p-value = 0.0069**)<br>IL-6: 2D vs Gel (p-value = 0.0924); 2D vs. TRACER (p-value = 0.0342*); Gel vs. TRACER (p-value = 0.857)<br>TNFα: 2D vs Gel (p-value = 0.1914); 2D vs. TRACER (p-value = 0.0045**); Gel vs. TRACER (p-value = 0.1516)<br><br>CCL22: 2D vs Gel (p-value = 0.9992); 2D vs. TRACER (p-value = 0.0018**); Gel vs. TRACER (p-value = 0.0017**)<br>CD23: 2D vs Gel (p-value = 0.314); 2D vs. TRACER (p-value = 0.0634); Gel vs. TRACER (p-value = 0.5869)<br>CD206: 2D vs Gel (p-value = 0.2809); 2D vs. TRACER (p-value = 0.4798); Gel vs. TRACER (p-value = 0.888) |
| **1B** | 3 independent donors, 3 technical replicates each | ANOVA with Tukey's multiple comparison post-hoc test | IL-1β, 2D: M0 vs M1 (p-value = 0.0697); M0 vs. M2 (p-value = 0.6532); M1 vs. M2 (p-value = 0.028*)<br>IL-1b, single-layer TRACER: M0 vs M1 (p-value = 0.0036**); M0 vs. M2 (p-value = 0.2692); M1 vs. M2 (p-value = 0.001***)<br><br>IL-6, 2D: M0 vs M1 (p-value = 0.022*); M0 vs. M2 (p-value = 0.9937); M1 vs. M2 (p-value = 0.034*)<br>IL-6, single-layer TRACER: M0 vs M1 (p-value = 0.1582); M0 vs. M2 (p-value = >0.999); M1 vs. M2 (p-value = 0.1871)<br><br>TNFα, 2D: M0 vs M1 (p-value = 0.0567); M0 vs. M2 (p-value = 0.8761); M1 vs. M2 (p-value = 0.1572)<br>TNFα, single-layer TRACER: M0 vs M1 (p-value = 0.02*); M0 vs. M2 (p-value = 0.9874); M1 vs. M2 (p-value = 0.0226*) |
| **1C** | 3 independent donors, 3 technical replicates each | ANOVA with Tukey's multiple comparison post-hoc test | CCL22, 2D: M0 vs M1 (p-value = 0.8827); M0 vs. M2 (p-value = 0.0432*); M1 vs. M2 (p-value = 0.091*)<br>CCL22, single-layer TRACER: M0 vs M1 (p-value = 0.8877); M0 vs. M2 (p-value = 0.0526); M1 vs. M2 (p-value = 0.0254*)<br><br>CD23, 2D: M0 vs M1 (p-value = 0.9989); M0 vs. M2 (p-value = 0.0022**); M1 vs. M2 (p-value = 0.0021**)<br>CD23, single-layer TRACER: M0 vs M1 (p-value = 0.9189); M0 vs. M2 (p-value = 0.0167*); M1 vs. M2 (p-value = 0.0262*) |
| **2A** | 2-3 biological replicates, 3 technical replicates each | ANOVA with Dunnett's multiple comparisons test, comparing against 100% CGM (no switch) | Day 0: vs. 0:100 blend (p-value >0.999); vs. 25:75 (p-value >0.999); vs.50:50 (p-value > 0.999); vs. 75:25 (p-value >0.999); vs. 100:0 (p-value > 0.999)<br><br>Day 2: vs. 0:100 blend (p-value =0.9998); vs. 25:75 (p-value =0.9999); vs.50:50 (p-value =0.9999); vs. 75:25 (p-value >0.999); vs. 100:0 (p-value > 0.999)<br><br>Day 4: vs. 0:100 blend (p-value =0.9755); vs. 25:75 (p-value =0.9458); vs.50:50 (p-value =0.9982); vs. 75:25 (p-value >0.999); vs. 100:0 (p-value > 0.9997)<br><br>Day 6: vs. 0:100 blend (p-value =0.1436); vs. 25:75 (p-value =0.1988); vs.50:50 (p-value =0.8185); vs. 75:25 (p-value =0.9785); vs. 100:0 (p-value = 0.9351)<br><br>Day 8: vs. 0:100 blend (p-value =0.0039**; vs. 25:75 (p-value =0.070); vs.50:50 (p-value =0.2393); vs. 75:25 (p-value =0.9611); vs. 100:0 (p-value = 0.9979)<br><br>*Ratio of media blend is organoid CGM: RPMI-1640* |

| | | | |
|---|---|---|---|
| **2B** | 2 biological replicates, 2-3 technical replicates each | ANOVA with Dunnett's multiple comparisons test | 0:100 (100%RPMI): Day 0 vs. Day 1 (p-value = 0.5941); Day 0 vs. Day 2 (p-value = 0.4985)<br>25:75 Day 0 vs. Day 1 (p-value = 0.3222); Day 0 vs. Day 2 (p-value = 0.3282)<br>50:50 Day 0 vs. Day 1 (p-value = 0.9471); Day 0 vs. Day 2 (p-value = 0.4077)<br>75:25 Day 0 vs. Day 1 (p-value = 0.9845); Day 0 vs. Day 2 (p-value = 0.9495)<br>100:0 (100% CGM) Day 0 vs. Day 1 (p-value = 0.0026**); Day 0 vs. Day 2 (p-value = 0.2731) |
| **2F** | 3 independent donors, 2-3 technical replicates each | Unpaired t-test comparing co-culture v. monoculture | IL-6 (macs), cc vs mc: 0h (p-value = 0.1544); 24h (p-value = 0.0181*); 48h (p-value = 0.2494)<br>VEGFA (macs), cc vs. mc: 0h (p-value = 0.1101); 24h (p-value = 0.0490*); 48h (p-value = 0.0761) |
| **2G** | 3 biological replicates, 2-3 technical replicates each | Unpaired t-test comparing co-culture v. monoculture | CSF-1 (ODCs), cc vs. mc: 0h (p-value = 0.1549); 24h (p-value < 0.0001****); 48h (p-value = 0.1387)<br>PD-L1 (ODCs), cc vs. mc: 0h (p-value = 0.0867); 24h (p-value =0.0842); 48h (p-value = 0.0150*) |
| **3B** | 3 biological replicates, 2-3 technical replicates each | Linear regression on L1 to L6 (Slope of regression line vs. 0) | Is slope significantly non-zero?<br>ODC MC (Yes, slope =; P-value = 0.0007 ***)<br>ODC CC (Yes, slope = ;P-value < 0.0001****)<br>Macrophage CC (Yes, slope = ;P-value = 0.007**) |
| **3C** | 3-4 biological replicates, 2-3 technical replicates each | Linear regression on L1 to L6 (Slope of regression line vs. 0) | Is slope significantly non-zero?<br>CHOP: ODC MC (Yes, slope = 3.026, p-value = 0.0011**); ODC CC (Yes, slope = 0.7227, p-value <0.0001****)<br>VEGFA: ODC MC (Yes, slope = 0.5129, p-value = 0.0254*); ODC CC (Yes, slope = 0.866, p-value = 0.0238*) |
| **3D** | 2-3 independent donors, combined 2 tissues for each donor (to obtain more RNA) | Linear regression on L1 to L6 (Slope of regression line vs. 0) | Is slope significantly non-zero?<br>GLUT1: Macrophage CC (Yes, slope = 2.416, p-value = 0.0047**)<br>REDD1: Macrophage CC (Yes, slope = 1.326, p-value = 0.0469*)<br>VEGFA: Macrophage CC (No, slope = 0.2439, p-value = 0.862) |
| **3E** | 3-4 biological replicates, 2-3 technical replicates each | Linear regression on L1 to L6 (Slope of regression line vs. 0) | Is slope significantly non-zero?<br>TGFβ: ODC MC (No, slope = 0.05378, p-value = 0.6069); ODC CC (Yes, slope = 0.2308, p-value =0.0279*)<br>PD-L1: ODC MC (No, slope = -0.04221, p-value = 0.3935); ODC CC (Yes, slope = -0.09151, p-value = 0.0267*) |
| **3F** | 3 independent donors, combined 2 tissues for each donor (to obtain more RNA) | Linear regression on L1 to L6 (Slope of regression line vs. 0) | Is slope significantly non-zero?<br>PD-L1: Macrophage CC (Yes, slope = 0.2765, p-value = 0.0436*) |
| **4D** | 3 biological replicates, 3 TRACERs each | Linear regression on L1 to L6 (Slope of regression line vs. 0) | Is slope significantly non-zero?<br>ODC MC (No, slope = 1.160, p-value = 0.5283)<br>ODC CC (Yes, slope = 4.363, p-value = 0.0244*) |
| **5B** | 3 independent donors treated with CM from 3 different TRACERs | ANOVA with Dunnett's multiple comparison test, against L1 | TNFα+: Mac MC vs. L3 (p-value = 0.9855); vs. L6 (p-value = 0.5111); vs. 0.2% hypoxia (p-value = 0.8205)<br>ODC MC: vs. L3 (p-value = 0.9994); vs. L6 (p-value = 0.6579); vs. 0.2% hypoxia (p-value = 0.9982)<br>CC: vs. L3 (p-value = 0.1852); vs. L6 (p-value = 0.0145*); vs. 0.2% hypoxia (p-value = 0.0185*)<br><br>GZMB+: Mac MC vs. L3 (p-value = 0.9877); vs. L6 (p-value = 0.37); vs. 0.2% hypoxia (p-value = 0.4761)<br>ODC MC: vs. L3 (p-value = 0.952); vs. L6 (p-value = 0.9838); vs. 0.2% hypoxia (p-value = 0.9965)<br>CC: vs. L3 (p-value = 0.5377); vs. L6 (p-value = 0.0454*); vs. 0.2% hypoxia (p-value = 0.0003***) |

| | | | |
|---|---|---|---|
| | | | TNFα+GZMB+: Mac MC vs. L3 (p-value = 0.978); vs. L6 (p-value = 0.9894); vs. 0.2% hypoxia (p-value = 0.9966)<br>ODC MC: vs. L3 (p-value = 0.975); vs. L6 (p-value = 0.959); vs. 0.2% hypoxia (p-value = 0.9677)<br>CC: vs. L3 (p-value = 0.1517); vs. L6 (p-value = 0.009**); vs. 0.2% hypoxia (p-value = 0.0007***) |
| **5C** | 3 independent donors treated with CM from 3 different TRACERs | ANOVA with Dunnett's multiple comparison test, against L1 | TNFα+: Mac MC vs. L3 (p-value = 0.0852); vs. L6 (p-value = 0.2964); vs. 0.2% hypoxia (p-value = 0.2902)<br>ODC MC: vs. L3 (p-value = 0.194); vs. L6 (p-value = 0.214); vs. 0.2% hypoxia (p-value = 0.9408)<br>CC: vs. L3 (p-value = 0.0218*); vs. L6 (p-value = 0.0009***); vs. 0.2% hypoxia (p-value = 0.0018**)<br><br>GZMB+: Mac MC vs. L3 (p-value = 0.7762); vs. L6 (p-value = 0.0179*); vs. 0.2% hypoxia (p-value = 0.0273*)<br>ODC MC: vs. L3 (p-value = 0.9664); vs. L6 (p-value = 0.6169); vs. 0.2% hypoxia (p-value = 0.4969)<br>CC: vs. L3 (p-value = 0.7351); vs. L6 (p-value = 0.3038); vs. 0.2% hypoxia (p-value = 0.0262*)<br><br>TNFα+GZMB+: Mac MC vs. L3 (p-value = 0.4895); vs. L6 (p-value = 0.914); vs. 0.2% hypoxia (p-value = 0.7605)<br>ODC MC: vs. L3 (p-value = 0.0.5542); vs. L6 (p-value = 0.8257); vs. 0.2% hypoxia (p-value = 0.4551)<br>CC: vs. L3 (p-value = 0.6884); vs. L6 (p-value = 0.0433*); vs. 0.2% hypoxia (p-value = 0.0186*) |

**Table S2: Primers for qPCR Analysis**

| Gene (Human) | Forward Primer (5' to 3') | Reverse Primer (5' to 3') |
|---|---|---|
| **18S** | GTAACCCGTTGAACCCCATT | CCATCCAATCGGTAGTAGCG |
| **GAPDH** | CTCTCTGCTCCTCCTGTTCGAC | TGAGCGATGTGGCTCGGCT |
| **CCL22** | GCCAACATGGAAGACAGCG | TTAGCAACACCACGCCAGG |
| **CD23** | AGGTGTCCAGCGGCTTTGT | CTTCCATGTCGTCACAGGCA |
| **CD206** | CTGTGCATTCCCGTTCAAGTT | CTGCCCTCAAATTTCAATGGAC |
| **IL-1β** | TTCTTCGACACATGGGATAACG | TGGAGAACACCACTTGTTGCT |
| **GLUT1** | CTGCTCATCAACCGCAAC | CTTCTTCTCCCGCATCATCT |
| **IL-6** | ACTCACCTCTTCAGAACGAATTG | CCATCTTTGGAAGGTTCAGGTTG |
| **RPLP0** | GTGCTGATGGGCAAGAAC | AGGTCCTCCTTGGTGAAC |
| **LYZ** | ACTACAATGCTGGAGACAGAAGC | GCACAAGCTACAGCATCAGCGA |
| **GATA6** | GTGCCCAGACCACTTGCTAT | TGGAATTATTGCTATTACCAGAGC |
| **KRT17** | ATCCTGCTGGATGTGAAGACGC | TCCACAATGGTACGCACCTGAC |
| **S100A2** | TGCCAAGAGGGCGACAAGTTCA | AAGTCCACCTGCTGGTCACTGT |
| **VEGFA** | TTGCCTTGCTGCTCTACCTCCA | GATGGCAGTAGCTGCGCTGATA |
| **PD-L1** | TGCCGACTACAAGCGAATTACTG | CTGCTTGTCCAGATGACTTCGG |
| **CHOP** | GGAGCATCAGTCCCCCACTT | TGTGGGATTGAGGGTCACATC |
| **REDD1** | GGTCACTGAGCAGCTCGAA | CCTGGACAGCAGCAACAGT |
| **TGFβ** | GCAGAAGTTGGCATGGTAGC | CCCTGGACACCAACTATTGC |
| **CSF1R** | ATCCGGCTGAAAGTGCAGAA | GGATTGCGAGCTTGGTGTTG |
| **SPP1** | GCCGAGGTGATAGTGTGGTT | AACGGGGATCGGCCTTGTATG |

| | | |
|---|---|---|
| **WNT7B** | CCCCCTCCCTGGATCATGCACA | GCCACCACGGATGACAGTGCT |
| **C1QC** | AGGATGGGTACGACGGACTG | GTAAGCCGGGTTCTCCCTTC |
| **TNFα** | TCAGATCATCTTCTCGAACCCC | ATCTCTCAGCTCCACGCCAT |
